# European maize genomes unveil pan-genomic dynamics of repeats and genes

**DOI:** 10.1101/766444

**Authors:** G. Haberer, E. Bauer, N. Kamal, H. Gundlach, I. Fischer, M.A. Seidel, M. Spannagl, C. Marcon, A. Ruban, C. Urbany, A. Nemri, F. Hochholdinger, M. Ouzunova, A. Houben, C.-C. Schön, K.F.X. Mayer

## Abstract

The exceptional diversity of maize (*Zea mays*) is the backbone of modern heterotic patterns and hybrid breeding. Historically, US farmers exploited this variability to establish today’s highly productive Corn Belt inbred lines from blends of dent and flint germplasm pools. Here, we report high quality *de novo* genome sequences of the four European flint lines EP1, F7, DK105 and PE0075 assembled to pseudomolecules with scaffold N50 ranging between 6.1 to 10.4 Mb. Comparative analyses with the two US Corn Belt genomes B73 and PH207 elucidates the pronounced differences between both germplasm groups. While overall syntenic order and consolidated gene annotations reveal only moderate pan-genomic differences, whole genome alignments delineating the core and dispensable genome, and the analysis of repeat structures, heterochromatic knobs and orthologous long terminal repeat retrotransposons (LTRs) unveil the extreme dynamics of the maize genome. Haplotypes derived from core genome SNPs demonstrate the tessellation of modern maize resulting from a complex breeding history. The high quality genome sequences of the flint pool are a crucial complement to the maize pan-genome and provide an important tool to study maize improvement at a genome scale and to enhance modern hybrid breeding.

## Introduction

Since its domestication ~10,000 years ago by Native Americans, maize (*Zea mays ssp. mays*) has become one of the most important sources for human nutrition and animal feeding. Extensive variation in landraces and breeding germplasm like dent or flint corns underpin the enormous phenotypic and genetic diversity of maize [1–3]. Today, US hybrids produced from inbed lines of different heterotic groups – eg. Stiff-Stalk and non Stiff-Stalk [4], are highly productive and agriculturally important worldwide. These US Corn Belt dents resulted from crosses in the nineteenth century between Southern dent lines introduced to the United States from Mexico, and Northern flints (NF) which were the predominant germplasm in the pre-Columbian era grown by the American Indians [5, 6]. Phenotypes and genotypes of NFs constitute a highly distinct maize germplasm and possess rounded hard kernels while dent lines contain a softer endosperm texture that caves in during maturation forming dentate kernels. Historical records and genetic and molecular data strongly indicate that NFs originated from American Indian populations of southwestern America and the Great Plains [2, 6, 7]. Improvement by early farmers and adaptation to cooler climate and different photoperiods extended NF growth as north as southern Canada. After the colonization of the New World, maize was spread to Europe from both the Caribbean Islands and northeast US [8]. The early maturing and cold tolerant flints were key to a successful maize cultivation in temperate regions of Europe. In consequence, Northern flint germplasm still has a major contribution to modern European maize breeding material while Corn Belt dent genomes contain on average one quarter from their blending with NFs [2, 6].

The complete genome sequence of the dent line B73 represented a milestone in maize breeding, genetic and genomic research [9]. Applying the rapid progress in sequencing technologies, the B73 genome and gene models have been updated several times to reference quality [10]. On the other hand, very high diversity at both the sequence and genic level has been reported between maize inbred lines for targeted regions and large-scale comparisons [3, 11–14]. Hence, the B73 reference sequence captures only a portion of the maize pan-genome. To overcome these limitations, a draft genome of the Iodent line PH207 [15], and - more recently-reference sequences of Mo17 and W22, two additional US dent lines, have been released [16, 17]. Genome-wide comparisons confirmed the high dynamics of the maize genome emphasizing the need for several high-quality genome sequences to characterize and exploit the pan-genomic variability in maize. Worldwide, many hybrid breeding programs focus on dent germplasm, whereas breeding programs in cooler regions of Central Europe exploit heterotic effects between dent and flint lines. Several studies have shown a clear differentiation of the Northern American and Northern European flint germplasm from the rest of the world [18]. While reference quality sequences exist for several dent inbred lines, the flint pool is still underexploited. To date, only one fragmented draft assembly with an N50 ~13.9 kb of one flint line, F2, is available covering ~65% of the estimated genome size [19]. We generated *de novo* reference sequences for four flint inbred lines representing predominant ancestors of maize hybrid breeding in Central Europe [20]. The four lines represent distinct flint germplasm sources such as populations Lacaune (F7, Southern France), Lizargarate (EP1, Northern Spain), Gelber Badischer Landmais (DK105, Southern Germany) and Petkuser Ferdinand Rot (PE0075, Northern Germany). Understanding the extent and quality of genomic differences between the flint and dent germplasm pools may contribute to a better understanding of the maize pan-genome and of the complementarity in heterotic patterns exploited in hybrid breeding. For a first assessment of the pan-genome, we contrasted gene and repeat contents of European flints to two US Corn Belt dents applying a consolidated gene annotation and whole genome alignments. The length and quality of the assemblies now allow aligning and comparing also repeat-dense intergenic regions and reveal the degree of repeat conservation and rapid turnover. Our analysis reveals the presence of widespread knob repeat “seed” regions and their influence on the surrounding gene landscape and expression characteristics. Finally, a haplotype-informed expression analysis surveys the effect of germplasm-specific regions on the maize transcriptome and reveals gene loci that are prime candidates for differences in the kernel phenotype among both gene pools. We conclude that for larger and complex plant genomes pan-genomic analysis will require a multidirectional angle and, beside gene-centric analyses, has to consider the structural and functional environment that might impose important *cis* regulatory effects.

## Results

### Whole genome sequencing

To access the genomic landscape of European flint maize, and to compare it with lines from the dent pool, we assembled four flint lines to reference genome quality. Three lines - EP1, F7 and DK105, are important founders of European breeding programs. The fourth line, PE0075, is a doubled haploid line derived from the Petkuser Ferdinand Rot population, a European landrace. We generated Illumina paired-end and mate pair sequences equivalent to ~ 220-320 x coverage (Supplementary Table 1), and assembled the reads to scaffolds and pseudo-molecules using the DeNovoMagic pipeline [15, 21, 22]. The large majority (94.3% up to 97.3%) of contig and scaffold sequences were integrated into 10 pseudochromosomes underpinning the high quality of the four new maize flint references (Table 1; Supplementary Table 2; Supplementary Figure 2A). We evaluated the completeness and sequence qualities of the four assemblies applying a BUSCO analysis [23]. Only ~2.5% of BUSCO genes were absent in each of the four assemblies on average, and another ~2% additional fragmented matches were observed. Complete BUSCO genes totaled >95% strongly supporting the high quality of the flint assemblies (Supplementary Table 2). Short read mapping of EP1 and F7 Illumina paired-end data supported a high sequence accuracy with less than one erroneous base per 100 kb (Supplementary Table 5). A genetic map generated from a F_2_ mapping population of a EP1xPH207 cross demonstrated a high consensus between genetic and physical map further corroborating the high quality and contiguity of the maize assemblies (Supplementary Figure 1).

**Table 1.** Genome statistics of maize dent and flint lines used in this study. Table summarizes total assembly size, total size of assembled chromosomes (^a^ only 10 pseudo-chromosomes), number of pseudo-chromosomes including unanchored scaffolds, N50 of scaffolds before pseudo-chromosome generation, total size of undefined sequence in the genome, number of genes (^b^ referring to the gene sets, see main text and supplement), syntelogs (referring to genes syntenic to at least one line), orthologs (i.e. syntelogs complemented by bidirectional best Blast hits for out-of-syntenic order genes) and total percentages of class I and class II repeats.

|  | *Zea mays* “flint” lines |  |  |  | *Zea mays* “dent” lines |  |
| --- | --- | --- | --- | --- | --- | --- |
| Line | EP1 | F7 | DK105 | PE0075 | B73v4 | PH207 |
| assembly size [Mb] | 2455.3 | 2392.8 | 2288.2 | 2198.5 | 2135.1 | 2156.2 |
| chromosome [Mb] <sup>a</sup> | 2321.0 | 2255.5 | 2176.1 | 2140.7 | 2106.3 | 2060.3 |
| scaffolds [#] | 60567 | 62610 | 797 | 972 | 267 | 43291 |
| scaffold N50 [Mb] | 6.13 | 9.48 | 10.39 | 8.64 | n.a. | n.a. |
| Undefined/gaps [Mb] | 25.48 | 25.62 | 14.33 | 14.67 | 30.73 | 442.21 |
| Genes, GS1 [#] <sup>b</sup> | 44,958 | 47,204 | 46,433 | 46,482 | 44,588 | 44,321 |
| Syntelogs GS1 | 36,479 | 36,811 | 36,729 | 36,801 | 37,274 | 35,158 |
| Genes, GS2 [#] <sup>b</sup> | 43,353 | 44,413 | 42,683 | 42,608 | 42,509 | 40,782 |
| Syntelogs GS2 | 38,847 | 39,126 | 39,058 | 39,208 | 39,526 | 37,726 |
| Orthologs GS2 | 40,308 | 40,483 | 40,560 | 40,621 | 40,753 | 39,253 |
| Class I repeats [% of assembly] | 79,1 | 78,6 | 78,3 | 77,8 | 77,2 | 74,9 |
| Class II repeats [% of assembly] | 2,03 | 2,07 | 2,11 | 2,16 | 2,28 | 2,29 |

### Pan-gene variation among flint and dent is moderate

To assess genic presence-absence variations (PAVs) in the six maize lines we employed a three-layer gene prediction pipeline. This ensured identification of gene variability among these lines and at the same time improved robustness of the structural gene annotation using comparative genomics components. For F7 and EP1 we predicted protein-coding genes as consensus models using protein homologies of known monocotyledonous protein sequences and a broad spectrum of transcriptome evidences from F7 and EP1 RNA-seq data (Supplementary Table 3). Initial gene sets comprised 39,352 and 41,387 protein coding genes for EP1 and F7, respectively. To exploit the potential of comparative genomics and to ensure a uniform and comparable gene annotation, we cross-mapped all gene models of the dent and flint maize lines and, eventually, complemented existing annotations. After filtering for transposon or transposon-derived genes this gene set (GS1) comprised ~45-47.5k protein coding gene calls per line (Table 1).

We evaluated and scored gene models by two metrics, orthologous relations and a multi-layer perceptron classifying genes into low (LC) and high confidence (HC) models, using the consistency of protein structures to known annotations. The eligibility of our classifier to identify problematic gene models is demonstrated by significantly lower expression levels of LC genes (median tpm_LC_ = 0.07 versus tpm_HC_ = 6.39), lower exon number per gene (2.9 exons per LC gene versus 5.3 exons per HC gene) and a higher fraction of single exon genes (29.1% versus 21.7%; Fisher test p < 1^−300^). In addition, out of 3,917 initial genes containing InterPro signatures indicative for transposons, 93% were classified as LC suggesting that the classifier is well capable to identify transposon-derived genes. The majority (~85-90%) of novel models added to GS1 were LC genes. However, between 650 and 900 HC genes complemented the initial structural gene calls per line. The second round of cross-consolidation limited to HC genes and LC genes with at least one syntenic counterpart adjusted gene structures and finalized gene set #2 (GS2) to a total of 41 to 44.5k genes with a mean of 42.5k genes per line. Variability in gene numbers is mainly reflecting underlying variations in assembly quality of the different lines. For example, the lower gene count found in PH207 is likely attributable to comparably larger gaps in the genome assembly whereas the higher gene count in EP1 and F7 is due to a higher amount of genes located in unanchored scaffolds that presumably represent assembly duplications.

Overall, 37,700 to 39,500 genes (88% to 93% of all genes per line) have a syntenic counterpart in at least one of the other lines and 95% of the genes in each genome had an orthologous copy in at least one of the other maize lines. Notably, GS2 increased the number of syntelogs by 2,200 to 2,600 genes per line. The cross-mapped annotations also identified between 1,380 (B73) and 1,680 (PH207) genes that were out-of-syntenic order but detected as a top hit by a syntelog of another line. 91.3% of these top hits represented bi-directional best blast hits (bbhs) to at least one of their proposed syntelog clusters suggesting that these orthologous candidates translocated in the respective maize line or represent small-scale misassemblies [24]. The remaining 1,376 up to 3,801 singleton genes had no syntelog cluster assignment. In all six lines, singleton genes were highly enriched for tandemly repeated genes, and the majority (60% in B73, PH207, DK105 and PE0075) of singletons were tandem duplications of syntenic genes indicating tandem gene duplications as a major source for gene copy number variations.

In summary, approximately 95% of the genes in each genome had an orthologous copy in at least one other maize line. GS2 significantly improved the completeness of syntelog clusters with 30,109 clusters that have syntelogs for all six lines (25,316 found for GS1) (Figure 1A and B). The addition of out-of-synteny bbhs further increased the number to 32,295 complete clusters, similar to the number of syntenic genes reported in the pairwise comparisons of W22 and Mo17 to B73 [17, 22]. However, shared pairwise gene content of GS2 – including out-of-synteny bbhs - range between 36,466 (between EP1 and PH207) and 38,133 (between DK105 and PE0075) indicating considerably higher conservation between our maize lines than previously reported.

**Figure 1.**
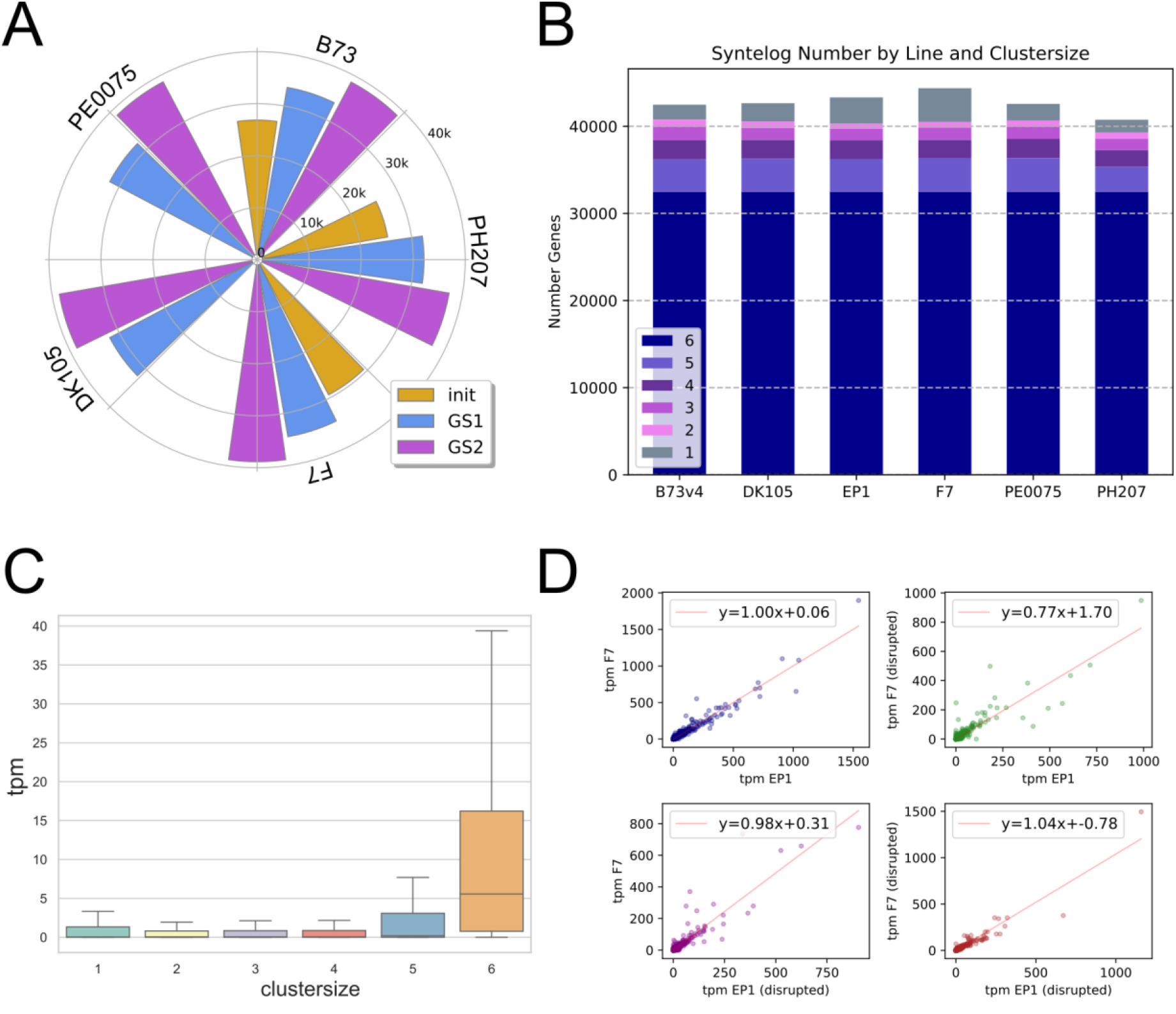
Gene consolidation and characteristics. **(A)** illustrates the iterative increase of pairwise syntelogs between EP1 and the other five maize lines applying an annotation consolidation by cross-mapped gene models. Syntelog count increases along the initial prediction (‘init’) and two consecutive consolidation steps reporting gene set 1 and 2 (GS1, GS2) for all pairwise comparisons. **(B)** shows the number of genes per line that are either singletons (label 1) or are syntenic between 2 and up to 6 lines (labels 2 to 6). **(C)** grades expression levels (tpm) of EP1 and F7 genes by size of their orthologous clusters. Genes either are singletons (cluster size 1) or have orthologues in one and up to five maize lines (cluster sizes 2 to 6). Syntelogs of orthologous clusters comprising 2-4 lines exhibit very low median expression levels (tpm ~0.06). **(D)** Consolidated mappings discovered several thousand genes for which the entire cross mapping of the top-scoring consolidation model had no contiguous but a disrupted ORF. These genes exhibit highly similar expression levels to their non-disrupted syntelogs (top right and bottom left panel). Overall, disrupted syntelogs have an equal linear correlation like non-disrupted syntenic pairs (top left panel).

Overall, core genes present in all maize lines exhibit high expression levels (mean and median 27.5 and 5.6 tpm) while PAV genes show significantly lower expression levels (median 0.07 tpm) and constitute the dispensable and highly variable gene fraction in maize (Figure 1C). The very low median expression of syntenic dispensable and singleton genes suggests no, only minor or highly specialized, line-specific functions for the majority of these genes.

Consistent with the genome analysis of the maize inbred line Mo17 [17], our cross-consolidation revealed a sizeable number of predicted genes with large effect mutations. Between 2,182 (B73) and up to 3,458 (PE0075) gene models had no contiguous open reading frame due to internal stop codons or frameshifts. Noteworthy, genes with disrupted ORFs had highly similar expression levels to their syntenic partner with contiguous reading frame. For example, pairwise comparison of F7 and EP1 syntelog expressions for the three classes – none, one or both syntelog(s) with a disrupted ORF – exhibited clear linear correlations with coefficients 0.8 < r < 0.95 (Figure 1D). Thus, a large majority of disrupted genes are not different in their expression to their syntenic partners with uninterrupted ORFs, and their mean and median expression are undiscernible to syntelog clusters from five lines. Such expressed a-, hypo- or even antimorphic alleles might represent a rich source of heterosis in maize and could also be involved in recently described nonsense induced transcriptional compensation (NITC) [25, 26].

### The full-length retrotransposon landscape illustrates the dynamics of transposable element (TE) generation and purging

Approximately 80% of the maize genome is constituted by repeats of different types. No pronounced differences in TE composition among the six lines analyzed was detected (Supplementary Table 6; Supplementary Figure 2). Full-length LTR retrotransposons (fl-LTR) have been shown to be very good indicators of the assembly quality and a linear relationship of the number of fl-LTRs and genome size has been observed [21]. For the maize flint lines sequenced and assembled in this study, we detected almost 15,000 high quality fl-LTRs per line, matching the expected genome size to fl-LTR ratio (Supplementary Table 7). The lower number observed for PH207 (6,838 fl-LTRs) can be attributed the overall lower quality assembly compared to the other five lines. Correspondingly, the PH207 assembly has to be treated with caution and depending on the analysis criteria was excluded from evaluation. For the remaining five lines, the number of fl-LTRs and their age distribution confirm the high quality of the respective assemblies (Supplementary Figure 3).

To quantify transposon dynamics between the different maize lines we identified still shared syntenic fl-LTRs by clustering TE junctions (50 bp outside and 50 bp inside the element) with high stringency. Only 8.9% of all fl-LTR locations where found to be shared between five lines. While the percentage of shared elements among the different lines shows a marked decrease, accordingly the pan-genome of the fl-LTR space almost doubles the number of elements. This high dynamic is in stark contrast to genes where 84.6% are retained at their syntenic position (Figure 2A). A pairwise cross comparison among the different genomes detected 23-31% of still syntenic fl-LTRs to be present in the corresponding line. The differences in pairwise shared numbers match the phylogenies from gene derived phylogenetic relationships (Figure 2B). Line unique fl-LTRs are younger and enriched in the more distal chromosome compartments concordant with higher recombination rates. Increasing the number of lines for shared fl-LTR elements reveals their continuous enrichment towards the distal low recombining compartments accompanied by an increase in age (Figure 2C). Our findings illustrate the rapid turnover of the intergenic space most likely driven by elimination through illegitimate recombination and, potentially, also functional consequences selected for by breeders and farmers during the cultivation and breeding process over the last centuries. The observation that lineage-specific transposable elements are more enriched in gene-rich regions towards the distal ends of the chromosomes, a genome environment that is usually lower in (fl-) TEs, and the observation that TE presence in the surrounding of genes frequently interferes with expression and functionality [27] thus allows to speculate that in part the lineage-specific TEs located in gene-rich regions might underlie functional and trait differences. Our results complement a similar recent comparison on the mobile genome content of four maize lines [28] and highlight the enormous dynamics of TE generation and purging which are already visible within the -in evolutionary time dimensions - very narrow time window after domestication and during the breeding process of maize and the spread along the Americas and towards Europe.

**Figure 2.**
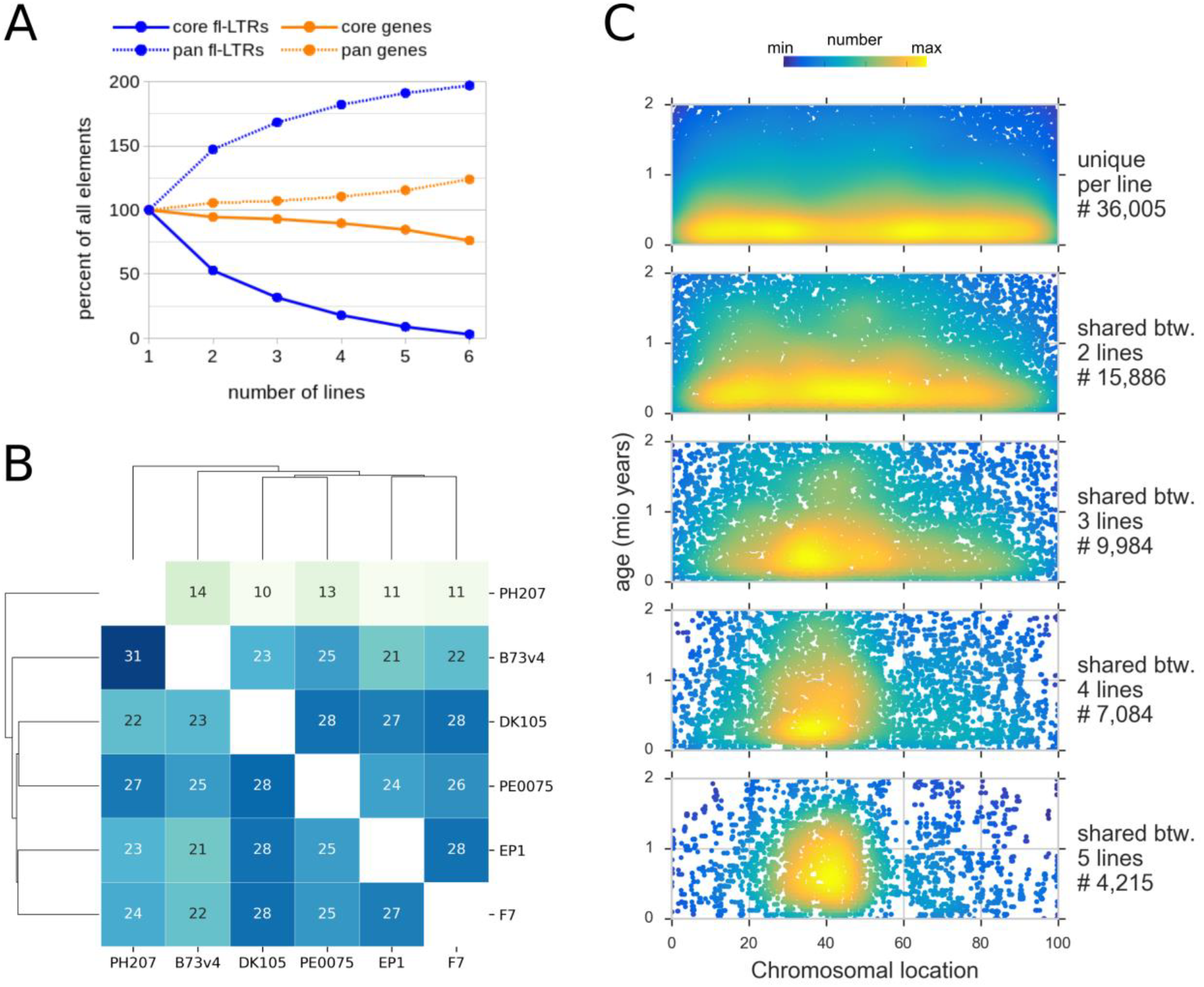
Pan and core characteristics of full length LTR-retrotransposons (fl-LTR). **(A)** Proportion of the pan and core sets for fl-LTRs and gene with increasing line numbers. **(B)** Percent of pairwise shared still intact fl-LTRs at syntenic positions for six maize lines. Reading direction is column to row, e.g. EP1 shares 27% of its fl-LTRs with F7 and vice versa F7 28% with EP1. The PH207 row has lower values because of its lower overall fl-LTR amount. Similar reciprocal values indicate a comparable assembly quality level. Based on the similarity matrix the six lines cluster into a relationship context, which distinguishes between flint and dent and places EP1 next to F7 and DK105 close to PE0075. A corresponding evaluation for genes gives pairwise shared numbers between 82 and 91 %. **(C)** Insertion age and chromosomal distribution of line unique and shared fl-LTRs in the different cluster constellations. The line unique fl-LTRs contain a higher proportion of younger elements. There is a continuous shift towards a more pericentromeric location and towards older elements if more lines share the same syntenic fl-LTR locus.

### Heterochromatic knob islands in the maize genome

Knob regions are heterochromatic regions in the genome that have been demonstrated to affect local recombination [29, 30]. They belong to the group of satellite tandem repeats which are well represented in our flint assemblies (Supplementary Figure 4 and 6A, Supplementary Table 6B). The Maize knobs are composed of 180 bp and closely related 202 bp tandemly repeated sequence units (Supplementary Figure 5C). Variations in position and extent of knob regions in maize have been documented [29, 31]. We compared knob positions and extent in the six lines analyzed using fluorescence in situ hybridization (FISH). A broad variety of position and strength of hybridization signals was observed (Figure 3A;). Intensity and position of knob regions detected for the different karyotypes clearly separate dent lines (B73 and PH207) from flint lines (EP1, F7, DK105, PE0075) (Figure 3A; Supplementary Figure 5B). In particular extensive knob regions detected on chromosomes 7 and 8 of the dent lines are not found in flint lines. For other positions on the genome, a clear flint/dent separation among lines cannot be observed. However, the variations observed among the lines reveals the pervasive dynamics of knob regions in the genome that also influence local recombination rates [29]. FISH only detects longer regions that carry the respective sequence signature while shorter regions containing fewer tandem units are not detectable. High quality genome assemblies resolving a larger proportion of the repetitive space now allow analyzing the so far hidden, cryptic, knob sites. We used the assembly of EP1, the best resolving assembly, to analyze for the characteristic sequence signatures. While all signals also detected by FISH (chromosomes 4L, 5L, 6S) have clearly detectable sequence counterparts in the assembly numerous additional positions with fewer tandem repeat units are found in the sequence assembly on all chromosomes (Figure 3B). Given the observations on the rapid diversification based on repeat units detailed above and the sequence and genome-based observations this might indicate numerous potential (shadowed) knob regions that can rapidly expand or shrink by e.g. illegitimate recombination and can cause recombinational isolation of the affected regions.

**Figure 3:**
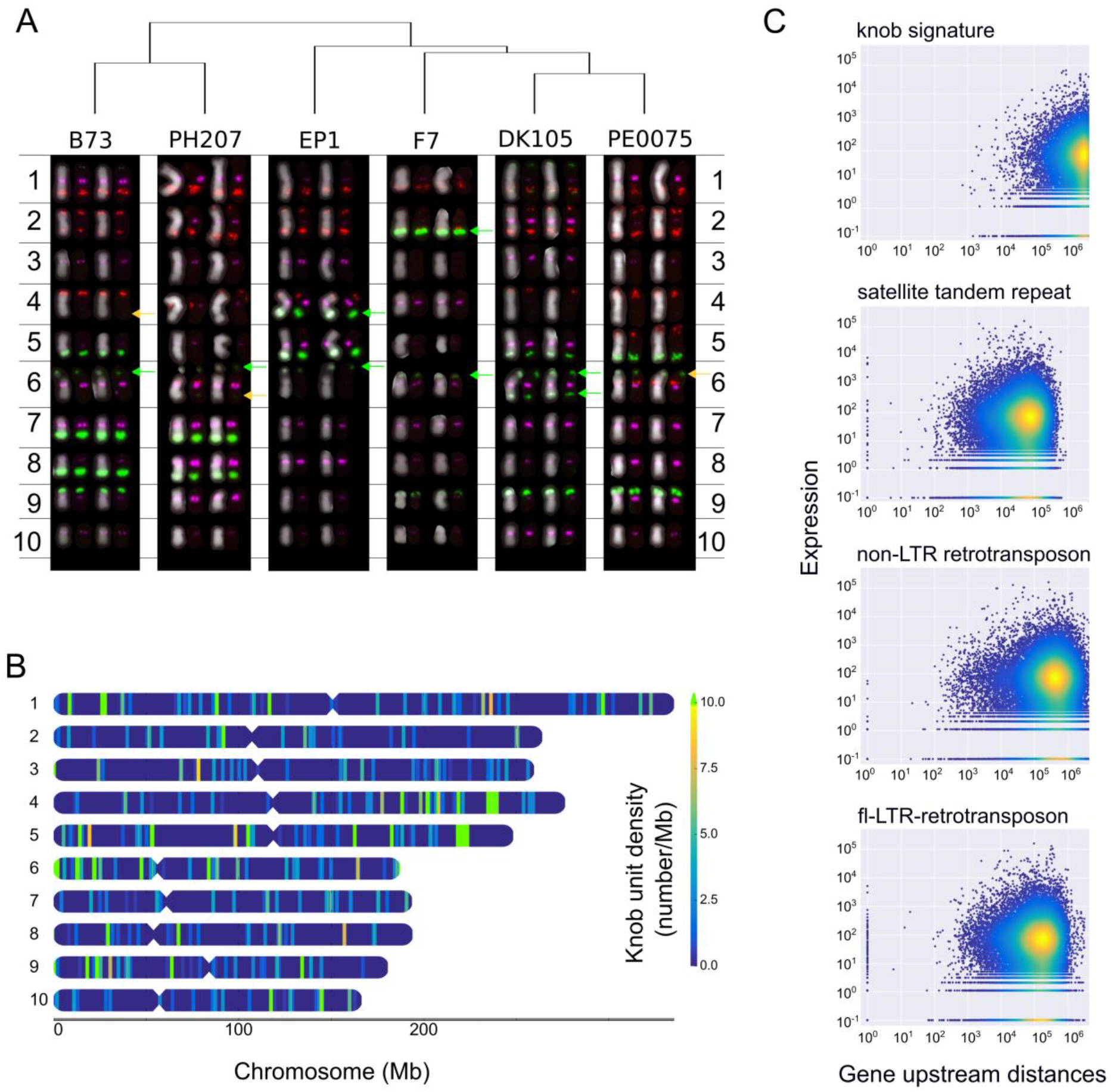
Diversity of knob locations in six maize lines. **(A)** Karyotyping of all six maize lines by FISH. Red-(ACT)10, green-Knob-2 (180 bp Knob repeat) and MR68-3 (Chromosome 6 clone MR68, ID AF020265.1), magenta-CentC69-1. Green arrows indicate co-occurrences of Knob-2 and MR68-3, orange arrows a sole occurrence of MR68-3. **(B)** Chromosomal locations of all knob sequences in EP1. Besides the three major Mb scaled knob regions identified by FISH on chromosome 4L, 5L and 6S many smaller knob sequences (<10 kb) are found scattered along all chromosomes. **(C)** Relation between gene neighborhood and gene expression. For each gene, the maximal expression of 7 different conditions is plotted against its upstream distance [bp] to the next neighboring element of a specific type. Both axes are logarithmic. It is noteworthy that especially knob sequences seem to be a “bad” neighborhood for genes. Here, in contrast to other element a 1,000 bp perimeter was found to be devoid of genes.

In fission yeast, knob structures have been shown to implicate significant downregulation of surrounding genes [32]. To check for similar effects in maize we analyzed the distance of genes to the nearest knob signature in the assembly and related it to expression data [33] generated for the lines B73, EP1 and F7. We find the close surrounding of knob repeat units to be devoid of genes (Figure 3C). The observation is even more pronounced than observed for other types of repeat elements and could be attributable to the heterochromatic characteristics of knob regions. Also for the genes surrounding knob regions, we observe a trend for lower expression values (Supplementary Figure 5D). Thus, the widespread presence of knob-type tandem repeats, the pronounced differences between lines, their rapid expansion and/or shrinking and the marked impact on the expression of genes located in the neighborhood might contribute to the appearance and impact of rare alleles and rare deleterious variants associated with crop fitness [33].

### Whole genome alignments

Numerous large structural variations (SV) in maize have been reported [11, 12, 14]. To gain insight into the extent of SVs in the maize flint versus dent genomes we generated whole-genome alignments (WGA) among all of the genomes used in this study. We thereby overcame limitations caused by commonly employed short read sequence read alignments (e.g. large extent of repetitive genome space with ambiguous mapping), and made use of the information gain provided by the full genome assemblies of all maize lines used in this study. Thus, contextual sequence and full genome information have been exploited and non-genic elements that make up for the overwhelming majority of the genomic space were considered for the WGAs.

We generated pairwise WGAs for the six lines and post-processed the resulting alignments to single alignment blocks (SAB) that represent the highest scoring one-to-one relationships between each genome pair. Single alignment blocks (SABs) were further concatenated to merged alignment blocks (MABs) if they followed a strict and unambiguous order in both genome sequences (Supplementary Figure 6). MABs provide an approximation of the overall contiguity between two genomes by bridging sequence gaps or regions of high sequence difference due to sequence insertions, deletions or inversions. On average, approximately 50% of the genome sequence aligned in each of the pairwise comparisons by SAB scoring(Supplementary Table 8). WGAs (MABs) assigned 80-90% (1.7 - 2 Gbp) of the genome sequences into 17,000 to 21,000 syntenic segments (Supplementary Table 9). Mean sizes of SABs and MABs were ~10kb and ~100 kb, respectively (Supplementary Figure 7). Consistent with previous findings, the analysis of MABs revealed numerous genomic rearrangements including translocations (~500 up to 1,650 inter-chromosomal) and inversions (~2,500 to 3,700) among the different genotypes. Inversions and translocations span a total of 48-115 Mbp and 5.3-12.8 Mbp of contiguous genomic sequences, respectively. While translocations are in general small and have a mean size of 8.6kb, the length distribution of inversions showed both small and larger sizes including 427 inversions ≥500 kb and up to 4.8 Mb (Supplementary Figure 8). All of them have potential influence on local recombination rates and separate individual lines genetically and functionally. To delineate regions that do not align with any other of the five lines (‘unaligned’), align solely with flint or dent maize lines (‘group-specific’) or are aligned in all pairwise alignments (‘core’), we combined pairwise WGAs of each line and projected them onto its genome sequence (Supplementary Figure 9). We classified a genomic region as core type if it aligned to four out of the five possible lines due to the high amount of gap sequences in PH207 that likely result in an underestimate of the true core genome. The core genome defined by SABs comprised on average ~850 Mb (40%) of the total genome while 71 Mb (3.3%) were group-specific and ~460 Mb (22%) unaligned. This indicates a large fraction of uniquely inserted or deleted sequences in each line. Positional analysis of genomic bins revealed significant positive correlations (τ ≥ 0.38; p<10^−36^) between the core and repeat density while gene densities were negatively correlated (−0.4 < τ < −0.33; p<10^−26^) (Figure 1A and B). Notably, for all six lines the highest core densities located at regions adjacent to the centromere corresponding to above average SAB sizes and below average recombination rates (Supplementary Figure 10). These observations are reminiscent of observations in the *Triticeae* gene space [21] mirroring the recombinogenic and functional properties of more central chromosomal regions vs. regions with higher recombination and genic innovation near the telomeric ends, characteristic for the dispensable genome. Recombination inertness of the central regions is also mirrored by the high repetitive content of these regions but also an overall better alignability compared to the more distal regions of the chromosomes.

### Genetic Mosaicism and Haplotypes in US Corn Belt dents and European flints

The core genomic regions served as positional anchors to generate multiple sequence alignments. One-to-one unambiguous (i.e. no overlapping tandem duplications) alignments for the six lines total 287 Mb (~15%) of each genome and comprise 6.25×10^6^ orthologous SNPs with positional information for each of the six genomic coordinate systems. The detected SNPs showed an excellent agreement of 99.3% (4,827 mismatches out of 653,398 total scored calls) to a published set of SNP calls using the 600K Affymetrix™ array [34]. This congruence is well within the error margins of genotyping arrays [35], and demonstrates the orthology and high accuracy of the multiple WGAs for the six lines. Given the observed SNP frequencies, and applying randomization studies and run-of-head statistics, the expected maximal run length of identical SNPs between two lines range between 21 to 30 consecutive SNPs. We applied a conservative threshold of ≥40 identical SNPs as seeds to identify shared haplotypes of likely common ancestry, and selected genomic windows with a minimum number ≥40 identical SNPs as seeds for a greedy extension to delineate genomic segments with ≥98% sequence identity between all 15 pairwise line combinations (Figure 4C). Mean and total genomic spans covered by these pairwise near-identical haplotypes range between 40 to 68 kb and 394 up to 664 Mb suggesting between one fifth and up to one third of the genome shared by a recent common ancestor. The proportion of shared genomic regions between flint and dent lines is in close agreement with previous studies of Northern flint content in Stiff-Stalk and non-Stiff Stalk dents [2, 6]. Number, mean and total sizes were highly consistent with their phylogenetic relationship deduced from high confidence orthologs (Supplementary Figure 11). However, each line of one group (flint or dent) contained substantial shared genomic portions of the other germplasm group (394 Mb up to 480 Mb for B73 and EP1 or PE0075, respectively). This is in line with the history of US Corn Belt dents that originated from crosses of Southern Dents and Flints of Northern America and Canada [6]. The latter established many founder lines of modern European flint breeding and were introduced to Europe starting as early as the discovery of the New World [8]. Alternatively, these haplotypes could be introgressions of US germplasm into European flint material that started in the 1950’s to broaden genetic diversity in European breeding programs. Notably, the DH line of the European landrace Petkuser exhibited the largest genome wide similarity of all flints towards B73.

**Figure 4.**
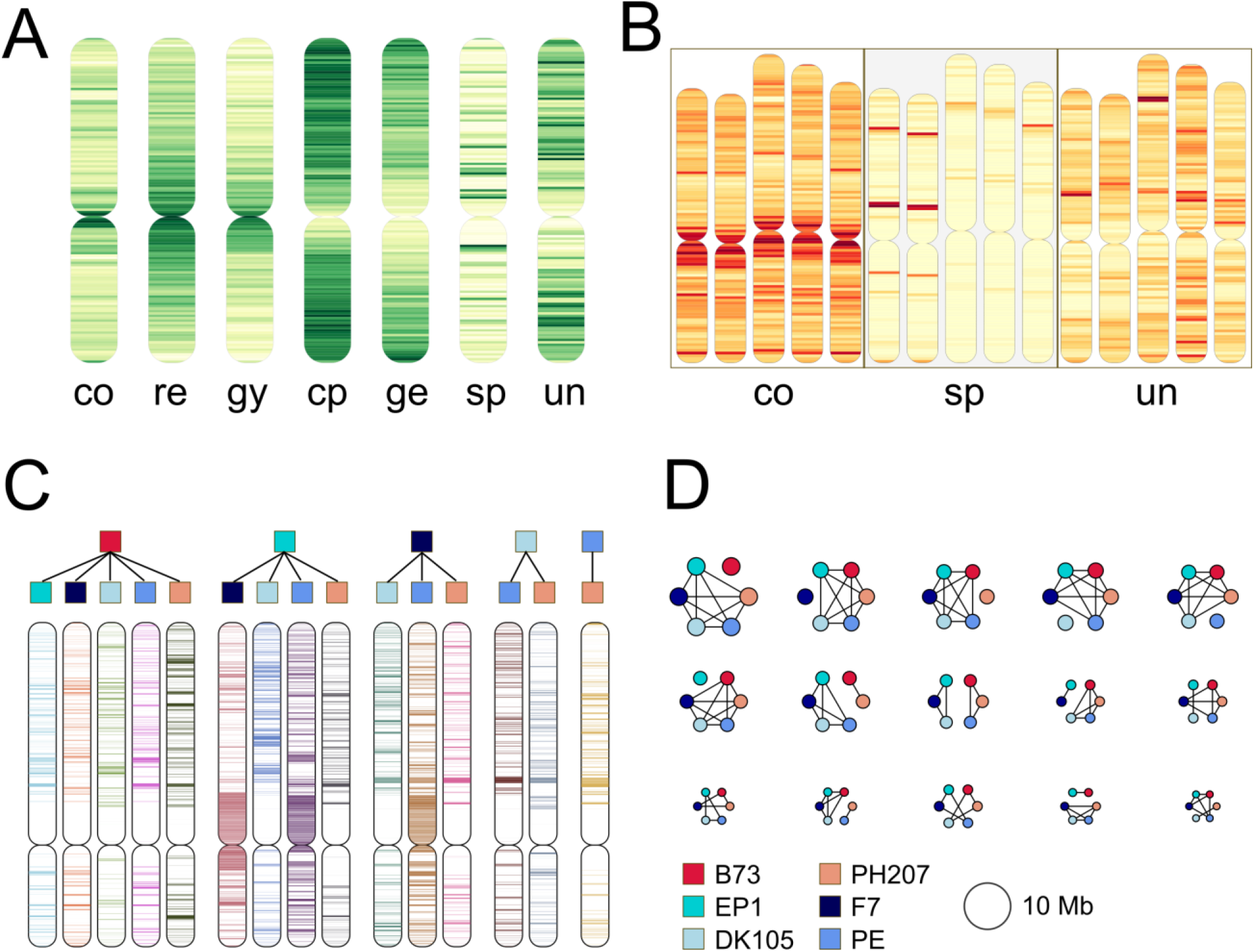
Comparative genome characteristics for the six maize lines. **(A)** shows the density distribution of the WGA (co: core, sp: germplasm-specific and un: unaligned regions) and functional genomic elements (re: all repeats, gy and cp: gypsy and copia LTR elements, ge: genes). The core WGAs are significantly positively correlated to the repeat and gypsy densities, similar to densities of copia elements and genes. For the unaligned and germplasm-specific densities, we detected no highly significant correlations with one of the other functional element. **(B)** Densities of the core, germplasm-specific and unaligned WGA regions exhibited similar distributions for all maize lines, exemplified for chr1 of B73, PH207, EP1, F7 and PE0075. Core regions are enriched at (peri-)centromeric regions, germplasm-specific obviously cluster in a group-specific manner and unaligned parts are random within all six lines. **(C)** shows runs-of-identity for SNPs from core regions for chr7 and all 15 pairwise combinations. Color code above the chromosome pictograms follows legend in 4D. **(D)** Sizes of higher order haplotypes of the six lines. Pictograms are proportional to their total genomic span, the legend provides the color coding of each of the six. Black lines indicate identity. Interestingly, the top six higher order haplotypes each group one maize variety as outlier and five lines that are identical by their SNP runs for these genomic segments. Of the more complex groupings, the most prominent haplotype separates the two germplasm, flint and dent.

Next, to pairwise runs-of-identities, we surveyed higher order haplotypes considering all 15 pairwise similarities simultaneously as combined binary patterns. Limiting these haplotypes to a minimum run size of 40 SNPs (see Methods), we identified 31 fitting our minimal size criterion (Figure 4D; Supplementary Table 10). Since the identified regions exceed random expectation by far and exhibit near complete SNP identity across particular line subsets/combinations, these regions are likely identical by descent and share a common ancestor. In total, the 31 distinct higher order haplotypes span ~288 Mb and comprise >1.1×10^6^ orthologous SNPs. Noteworthy, ordering by the number of SNPs as well as the genomic region covered, the top five/six haplotypes encode combinatorial groupings with five out of six lines being identical. The haplotype distinguishing between the studied dent (B73, PH207) and European flint lines ranked only as the sixth (by size) or seventh (by SNP number) most frequent (Figure 4D; Supplementary Table 10). Pairwise and higher order haplotypes depict the high genetic mosaicism of the pan-genome built from European flint and the two US Corn Belt dent lines. The distinct higher order haplotypes distribute quite evenly along the 10 chromosomes. We observed, however, also several striking clusters: for example a cluster on chromosome 3 with the three flints as one group and the 2 dents together with the Petkuser line form two distinct IBD regions. For two other regions on chromosome 2, five lines show identical SNP patterns while either F7 or B73 had distinct haplotypes. Though not the most prominent IBD type, three regions were enriched for the haplotype distinctive between flint and dent lines: two at the distal sites of chromosomes 4 and 7, and a large portion spanning a total around 12 Mbp in the proximal part of chromosome 8. The germplasm specific haplotypes differentiating our flint and dent lines strongly coincide with the group-specific WGAs on chromosome 8 (Supplementary Figure10) and enclose the major flowering time QTL *vgt1* in maize for which a *cis*-element 70kb upstream of the AP2 transcription factor gene rap2.7 has been proposed as causative locus [36, 37].

### Expression differences among different maize lineages

To explore putative functional consequences of the haplotype clearly differentiating our dent and European flint lines, we analyzed its effect on the maize transcriptome. From the maize association panel comprising 282 lines we selected a subset of 40 lines, which were highly similar to either B73 or F7/EP1 within regions identified by the haplotype analysis (Supplementary Figure 13) [38, 39]. We then analyzed genome-wide expression levels between the two contrasting groups using expression data from seven different conditions and tissues including root, shoot, leaf base and tip, leaf under light and dark, and kernels [33]. In total, 7,684 genes were differentially expressed (DEGs) over all seven tissues and conditions (Figure 5). A major phenotypic difference between the two germplasm groups analyzed is the kernel type. Maize endosperm contains soft starch granules that are embedded in a vitreous protein matrix of zein prolamines. Dent corn has softer starch at the kernel top, which caves in during maturation forming the characteristic tooth-like shape while flint kernels have a harder outer shell and remain round shaped.

**Figure 5.**
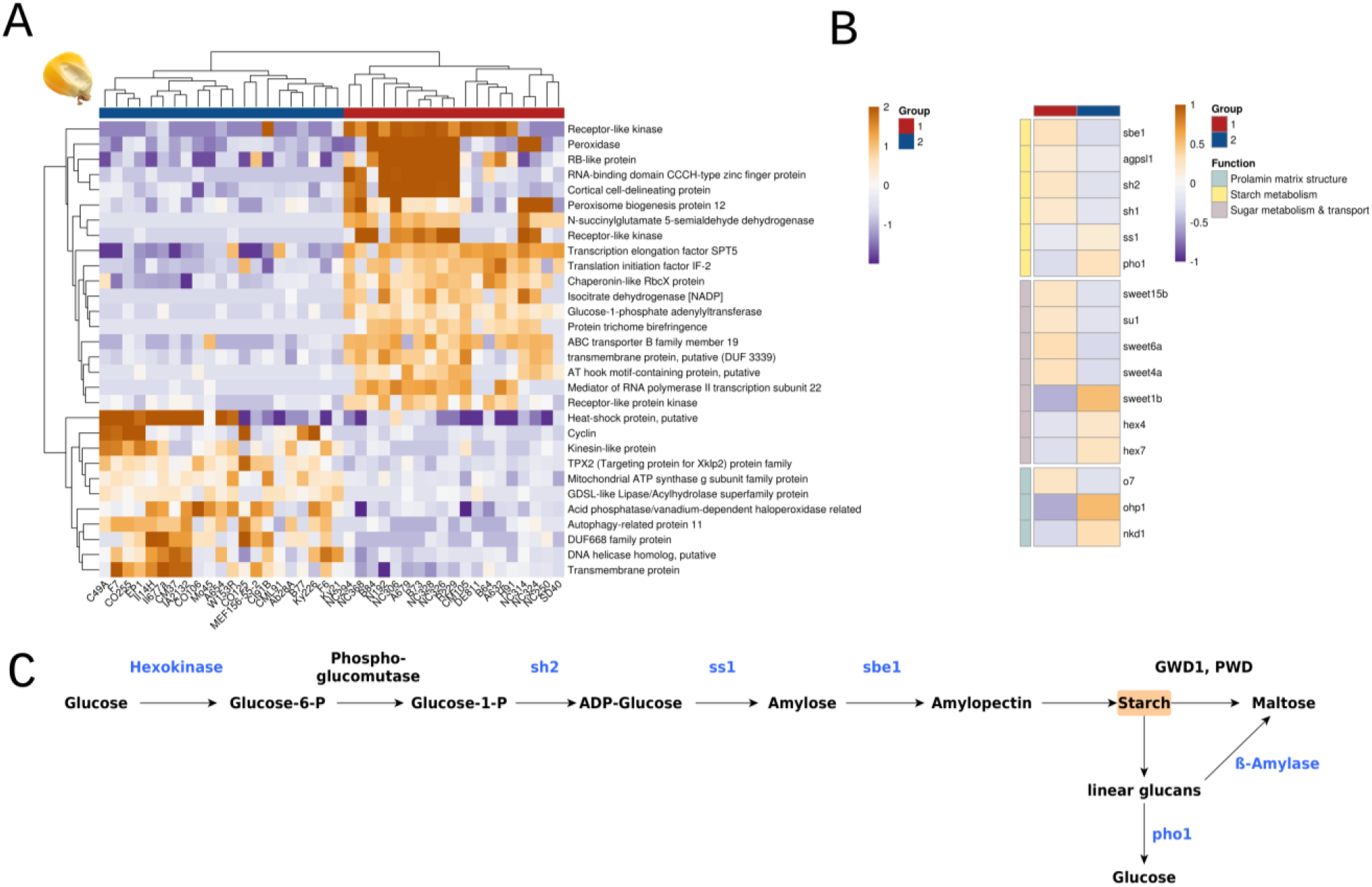
Haploblock-informed differentially expressed genes. **(A)** Heat map of top 30 differentially expressed genes (log2 fold change above 2 or below −2, adjusted p-value <= 0.05, variance stabilized) between the two haploblock-informed groups in kernels. Group 1: lines with haploblock-similarity to B73 (dent-alike), group 2: lines with haploblock-similarity to EP1/F7 (flint-alike). **(B)** Differentially expressed genes between the two groups that encode for components of the starch biosynthesis pathway, for proteins involved in sugar metabolism and transport into the kernel and for proteins organizing the prolamin matrix structure of the kernel. (sbe1: starch branching enzyme 1; agpsl1: ADP-glucose pyrophosphorylase small subunit leaf1; sh2: shrunken 2; sh1: shrunken 1; pho1: starch phosphorylase 1; sweet: sugar transporter; su1: sugary 1; hex: hexokinase; o7: opaque7; ohp1: opaque2; nkd1: naked endosperm 1). **(C)** Main reactions of starch biosynthesis and catalyzing enzymes, genes differentially expressed between the two groups of maize lines are indicated in blue color.

Scanning the identified DEGs for curated maize genes with known functions and phenotypes revealed a striking pattern for genes involved in the establishment of the kernel texture. We identified a plethora of genes involved in starch metabolism: sugar transporters including several proteins of the SWEET family, *hexokinase 4* and *7*, sucrose synthase *shrunken1*, the rate-limiting gene for starch biosynthesis (*shrunken2*), *starch synthase 1* (SS1), *starch phosphorylase*, the α(1→6)-glucosylhydrolase *sugary* [40] and starch branching enzyme *sbe1* catalyzing the formation of α(1→6)-glycosidic bonds in amylopectin (Figure 5C). Interestingly, the rice ortholog of *rap2.7* is *rice starch regulator 1* (*rsr1*) for which it has been shown that a T-DNA mutant has altered ratios of amylose to amylopectin and different starch granularity [41]. Carbohydrate metabolism complexly correlates with flowering time and *rap2.7* may therefore have a dual role in maize [42, 43]. In summary, this set of differentially expressed genes widely covers synthesis and turnover steps of the starch pathway.

We also detected a number of genes that function in the establishment and regulation of prolamin content and structure of the maize endosperm (Figure 5B). These genes encode the direct transcriptional regulator *ohp1* of α and γ zeins and several genes regulating the packaging of zeins and starch granules: opaque endosperm *o7* and *o10*, *empty pericarp 16*, floury endosperm *fl1* and naked endosperm *nkd1* [44–47].

## Discussion

Currently available maize reference genomes were limited to four US dent lines, B73, PH207, W22 and Mo17 [10, 15–17]. The full genome sequencing of the four different European flint genomes EP1, F7, DK105 and PE0075 to reference quality closes an important knowledge gap in maize research and complements the maize pan-genome. Gene and repeat content together with the analysis of WGAs and SNPs confirms the European flints as highly different germplasm of cultivated maize. For all these analyses, the four flint and the two dent lines, B73 and PH207, form two distinct groups, with each line sharing larger numbers of syntenic genes, repeats, aligned regions and SNP genotypes with members of the same group than with any member of the other group. Nevertheless, we also observed significant overall conservation of chromosome structure and gene content. Aligned blocks ordered by consecutive syntenic order span ~85% of the entire genome in all lines. A moderate number of non-syntenic genes also reflects this high degree of syntenic conservation. Our comparative cross-mapping strategy complemented in all lines a considerable number of previously missed genes. Consistent with reports in Mo17 [17], enforcing full-length mappings of top-scoring consolidation models revealed a substantial fraction (>8% of the annotations) of genes that contain large effect mutations and disrupted ORFs. These may contribute to widespread dominance effects and consequently heterosis reported in several studies in maize [48]. The consolidated annotation suggests on average 35-36,000 syntenic protein-coding genes and another 1,000-2,000 non-syntenic orthologues shared between two lines. Taking larger sequence gaps of PH207 into account, we estimate the core gene number in maize from the cluster sizes comprising syntelogs of five or six lines to range between 35-36,000 genes. A considerable number of non-syntenic genes were tandemly repeated genes suggesting a significant contribution to genic CNVs. Moreover, a large number of syntelogs found in only a subset of lines were expressed very lowly or not at all in a dataset of 26 different tissues and conditions contesting functionality of these predictions. However, the latter set also contains genes expressed at moderate to very high levels, which might contribute to line-specific adaptations and are of high interest for maize improvement and breeding. Additionally, some of these genes may have highly confined expression patterns or conditions that were missed in our transcriptome analysis. Though genic PAVs were moderate in our maize lines, there are also clear pieces of evidence that genic copy number variations play an important role in the differentiation of maize germplasm. Assessing the abundance of all four combinations of the set of syntelogs shows the largest difference for the division into 4 flint versus 2 dent lines (Supplementary Figure 12) clearly demonstrating that genic PAVs adhere to the expected pedigree information and germplasm relationships.

In contrast to the overall syntenic conservation, WGAs, the analysis of fl-LTRs and knob repeat patterns demonstrated enormous diversity and dynamics of the maize genome. This is also basis for the high amount of unaligned, non-orthologous genomic sequences, which are significantly larger than germplasm-specific WGAs and approximately ~½ of the core genome. Even more pronounced, however, is the asymmetry for shared orthologous fl-LTRs. Less than one tenth of them are conserved in all six lines. Consistent with previous reports, and similar to LTRs, large heterochromatic knobs display substantial variation in position (and likely also extent) among the maize lines studied [31, 49]. Remarkably, both shared sets of orthologous LTRs and knob positions perfectly reproduced the relationship of the six lines based on coding sequences, indicating smooth transitions between these features during breeding and admixture. In contrast to previous studies reporting low knob numbers in NF or even knob-less NF lines, the European flints including DK105 and the Petkuser line, contained similar knob numbers like the US dents [49]. It is unclear whether this is a characteristic of the NF ancestral lines migrated to Europe, reflects the breeding history in European flints or represents a technical advantage given the progress in methodology to detect weaker signals. For the first time, larger or nearly complete knobs - particularly for EP1-assembled into pseudomolecule sequences. Consistent with fiber-FISH studies, we also detected numerous locations of the 180 bp knob repeat unit throughout the chromosome assemblies [50]. These widespread knob repeats affected gene density and expression levels of adjacent genes adding further complexity to known effects of large-scale knobs in maize on recombination rates, meiotic drive and phenotypic traits like flowering time [29, 51, 52].

WGAs enabled to identify long runs of pairwise and higher-order combinations of genomic regions with identical haplotypes. Size and distribution of these haplotype blocks reflect complexíty of historic recombination, intercrossing and breeding history over the last centuries [3, 5, 7, 18, 53–55]. Similar to the gene and repeat comparisons, the SNP runs highlight the separation of flints and dents of this study but also illustrate and delineate candidate regions of common ancestry in the US Corn Belt and European flint lines. We identified a set of regions that differentiated the studied Corn Belt dents from the European germplasm. Intriguingly, genes differentially expressed between two groups of maize varieties that have been selected by their genotypic similarity to these regions comprised a nearly complete starch biosynthesis pathway and several genes involved in the establishment and organization of the endosperm texture. These loci are certainly good candidates for the genetic analysis of hard and soft kernel phenotype of flint and dents, respectively. However, within each group of our selected maize varieties, there are also lines classified as semi-dent or semi-flint. The complexity of the kernel/endosperm phenotype suggests that probably hundreds of loci contribute to it and that their impact will be on many regulatory levels, including transcriptional regulation but also other *trans*-active effects like protein-protein interactions or *cis*-effects from alleles modifying protein function. Hence, we speculate that altered expression of the identified genes represent a predisposition for the kernel phenotype rather than the sole causative factors. Moreover, the genetic mosacism observed within our six lines suggests that there are few if any such factors specific for one of the germplasm groups. Nevertheless, besides the detection of selective sweeps and other genomic signatures, it might be equally informative to survey *trans*-effects of such genomic regions. We demonstrate the value of full genome sequencing and assembly - rather than restriction to resequencing approaches -that allows analyzing genomic variations beyond the sole gene complement. While sequencing and assembly of complex grass genomes, such as maize, was for a long time complicated by technical and economic limitations, recent developments in assemblies of complex plant genome now appear to be reasonably economical and feasible. Haplotype characterization and contemporary methods to e.g. calculate breeding value and genome wide association mappings (GWAS) might profit from complementing gene-centric analysis by differences of knobs and LTRs in the circumference of genes associated to traits of interest.

## Supporting information

european_maize

## Acknowledgements

We thank Nina Opitz (University of Bonn) for helping to collect maize tissue for RNA-seq.

## Funding

This study was funded by the Bavarian State Ministry of the Environment and Consumer Protection (Grant ID TGC01GCUFuE69741; Project BayKlimaFit; http://www.bayklimafit.de/) and the German Federal Ministry of Education and Research (BMBF) within the funding initiative “Plant Breeding Research for the Bioeconomy” (Grant ID 031B0195A; Project MAZE; http://www.europeanmaize.net). In addition K.F.X Mayer acknowledges support from the BMBF within the project de.NBI Grant ID 031A536B.

## Author contributions

S.C-C., M.KFX., O.M. and H.G. designed the study. M.KFX. and H.G. wrote the manuscript. B.E. and S.C-C. supervised the genomic sequencing. B.E., U.C. and N.A. generated the F2 population and constructed the genetic map. H.G. analyzed genes and whole genome alignments. G.H. performed the repeat analysis. H.A. and R.A. contributed the FISH analysis. M.C. and H.F. prepared and sequenced the RNAseq samples. K.N., S.MA. and F.I. conducted the DEG expression analysis and the phylogenetic analysis. S.A. and S.MA. provided bioinformatics support. All authors critically read and edited the manuscript.

## Methods

### Plant material and Genome assembly

The four flint inbred lines were chosen to represent landraces from different European ancestry. While PE0075, a doubled-haploid line derived from the landrace Petkuser Ferdinand Rot, and DK105, derived from Gelber Badischer Landmais, can be classified as a representatives of the Northern flints, EP1 and F7 represent Pyrenean-Galician ancestry and were derived from populations Lizargarate and Lacaune, repectively [56]. EP1, F7 and DK105 are important founder line of European breeding programs. All four lines were assembled using the DeNovoMAGIC pipeline of NRGENE (Energin. R Technologies 2009 Ltd.; Ness Ziona, Israel). The application of this toolset has been previously described for several plant genome assemblies [15, 16, 57]. Briefly, genomic DNA was prepared from bulked leaf tissue from 15-24 seedlings per line. Leaf tissue was harvested, immediately ground in liquid nitrogen and DNA was isolated using a modified CTAB protocol [58]. Illumina sequencing of paired-end (PE) and mate-pair (MP) libraries was performed as specified in Supplementary Table 1. Contigs, scaffolds and pseudo-chromosomes were assembled *de novo* using the DeNovoMAGIC 2.0 technology for lines EP1, F7, and DeNovoMAGIC 3.0 for lines DK105 and PE0075, respectively, using the B73v4 reference genome to assist anchoring and orienting scaffolds and contigs to pseudo-chromosomes (Supplementary Table 2).

A cross of parental lines PH207 x EP1 was used to establish a high-density genetic map from an F_2_ mapping population. DNA from the parental lines and 192 F_2_ plants were analysed using the Affymetrix Axiom™ Maize 600k Array and data were processed as described by [34]. Genotype calls from the quality category “Poly High Resolution” were filtered for markers polymorphic between the parental lines, allowing no missing values, which resulted in 174,616 markers. This dataset was processed through a script from the POPSEQ pipeline to cluster markers into groups showing identical segregation patterns [59]. A first genetic map was calculated with 9404 binmap markers with the R package ASMap v. 1.0-2 [60] using the function mstmap with the following parameters: pop.type="RIL2", dist.fun="kosambi", objective.fun="COUNT", p.value=1e-22, noMap.dist=15, noMap.size=2, miss.thresh=0.00. Linkage groups were assigned to the ten maize chromosomes based on previously mapped markers. Unlinked small groups with only a few markers were discarded. Ten F2 plants exhibited very high numbers of crossovers and were excluded from further analyses. In multiple rounds of mapping using the same parameters as stated above, markers with highly distorted segregation or which led to double-crossovers were identified using the function statMark in the ASMap R package and discarded before final map construction. The final genetic linkage map contained 8869 markers. All markers from the initial dataset which had a Hamming distance of 0 with one of the mapped binmap markers were inserted into the map, resulting in a genetic map with 174,071 markers.

### Gene annotation, Consolidation and Synteny

We predicted protein-coding structures for F7 and EP1 as consensus models using an approach as previously described [61, 62]. Briefly, consensus gene models are based on transcriptome evidences from F7 and EP1 RNAseq data generated within this study (see RNA preparation and expression analysis) and protein homologies of known monocotyledonous protein sequences (maize B73 and PH207, *Sorghum bicolor* v3.1., *Oryza sativa* v7 MSU and *Brachypodium distachyon* v3.1) [61, 63, 64]. Evidences were mapped onto the genome assembly applying GenomeThreader [65] using a minimum alignment coverage of 50% and seed sizes between 7-10 for protein and 18 for nucleic acid matches. Before mapping, transcript data were assembled using Trinity with default parameters and Bridger with kmer sizes 25 and 29 bp [66, 67]. Subsequently, these sequences were combined by the evidential gene pipeline (http://arthropods.eugenes.org/EvidentialGene) to obtain the final EP1 and F7 transcriptome assemblies. Additional candidate loci for the initial gene sets of EP1 and F7, and the original representative gene models of B73 and PH207 [68] were searched in all six genome sequences (including PE0075 and DK105) by blat alignments [69]. We denote gene models used for the mapping as ‘informants’, and those derived from informant mappings to another genome sequence as ‘target’ models. Homologous loci were further refined by exonerate surveys [70] spanning the genomic region detected by blat and 20 kb up- and downstream sequence. Blat and exonerate alignments showed a linear correlation allowing direct conversion and comparison of both scoring schemes (r>0.999). For each blat/exonerate target pair, the top scoring model was selected and added to the initial models if the target (i) did not overlap with existing gene models, (ii) had a contiguous ORF, and (iii) ≥95% of the informant sequence was covered by the target. The resulting gene set constituted gene set 1 (GS1). Current gene annotations largely rely on automatic pipelines and still contain poor gene models like partial, merged or transposon-derived structures. Cross mapping bears the danger to transfer such models between each genome and thereby potentially enriches poor annotations. To estimate this effect and to evaluate gene-based statements of this study, we trained a multi-layer perceptron (MLP) classifying genes into low (LC) and high confidence (HC) models based on their consistency to known protein structures of other annotations. We applied the implementation of scikit-learn (https://scikit-learn.org) of an MLPclassifier using backpropagation learning with the stochastic gradient-based optimizer, and two hidden layers of 6 and 3 neurons, respectively. A training set was based on swissprot maize proteins and a set COG clusters [71] that were constructed from bi-directional best blast hits (bbhs) between 10 plant species: *Arabidopsis thaliana, Oryza sativa, Hordeum vulgare, Triticum aestivum, Sorghum bicolor, Sertaria italica, Brachypodium distachyon, Ananas comosus, Leersia oryzoides, Phyllostachus edulis*. At least three mutual bbhs with an alignment coverage ≥95% defined a COG. The high confidence (HC) training set (‘positives’) consisted of maize swissprot proteins entirely matching a maize fl-cDNA

[72] or B73 proteins with an alignment coverage ≥95% to one of the COG clusters above. Maize genes with matches <50% coverage to any of the COG cluster genes formed the low confidence (LC) set (‘negatives’). For each gene in both sets, we constructed an 18-tuple feature vector recording identity, query and hit coverage of its top blastp match against Arabidopsis, Rice, Sorghum, Brachypodium, emmer and all curated swissprot proteins. Positives and negatives were well separated by PCA, and 10x cross-validation showed a high accuracy >99% of the MLPclassifier. A set of 3,917 transposon genes identified from GS1 based on Interpro domains and description lines (https://github.com/groupschoof/AHRD) was excluded from training data and separately classified afterwards. It should be emphasized that LC genes cannot solely considered as biologically non-functional genes but comprise both, true genes displaying erroneous structures caused by problematic underlying evidences, sequences, gene calls etc., as well as potential over-predictions.

To derive gene set 2 (GS2), we first selected HC singletons and genes from syntelog clusters (see below) that comprised only syntelogs of maximal 5 lines and/or syntelogs differing from the CDS mean size of the respective cluster by > 5%. Each of these gene models were cross-mapped to the other five lines using the blat/exonerate alignment procedure described above. We collected all matches between the target and the informant genome that were located at a neighboring (±5 Mb) syntenic position as defined by the WGA (see below). If the informant model was located outside of an alignment, we approximated its syntenic position using WGAs framing the informant and set the midpoint in the target genome proportional to the distances to both flanking alignments. Subsequently, for each syntelog cluster we determined the top scoring combination of matches over all lines to infer gene models that optimize the fit and score for all participating lines and matches of this cluster. First, GS1 models and all selected matches were clustered into groups with congruent CDS sizes (maximal difference ±5%). For each group, we selected the top scoring model per line and summed their scores to obtain a group score. The final annotation was then based on models of the top scoring group and replaced or complemented by the GS1 model in one or more lines of the syntelog cluster. This gene set was further cleaned for transposon derived genes by the removal of singleton genes and syntelog clusters for which more than half of the cluster genes showed Interpro domains matching transposon domains.

We applied i-adhore v3 [73] running the hybrid cluster mode, a tandem gene distance of 10 and a minimum of 5 anchors to generate higher order syntenic relations for all six maize lines. Input pairwise gene similarities were derived from all-against-all blastn searches against coding sequences (CDS). To address putative ambiguities in the syntelog assignment caused by tandem arrays and the WGD in maize, we constructed a syntelog graph G = (V,E) with gene IDs as vertices V and multiplicon pairs provided by i-adhore as edges E. Edge weights were proportional to the size of a multiplicon, ie. the number of syntelogs within one syntenic block. In case of multiple edges from one vertex/gene, we only kept the highest scoring edge(s). This rule was applied in two consecutive steps, first to intra- and then inter-line connectors. In a second post-processing step, remaining clusters with vertices having two or more links to one line were further split into maximal cliques and a score was assigned to each clique as the sum of its edge/multiplicon weights. Next, we disjoint the cluster graph into non-overlapping clique subgraphs with a top-down approach thereby keeping cliques with lower scores only if none of its nodes is contained in the already selected subgraphs. The underlying rationale to select for syntelog cliques is analogous to the bidirectional best blast hit and the COG schemes.

### Fluorescent *in situ* hybridization (FISH) and karyotyping

Maize chromosomes were prepared from root meristems of two days old seedlings. Roots were cut and treated with 2mM 8-hydroxyquinoline solution for 3.5 h. Fixation was performed overnight at room temperature in 3: 1 (ethanol: acetic acid) fixative. Slides were prepared according to [74] and *in situ* hybridization was performed as described in [75]. Oligonucleotide probes for karyotyping and identification of knob repeats were chosen from [76]. Following sequences were synthetized and 5’-labeled by Eurofins Genomics (Ebersberg, Germany):

(ACT)_10_ : [TAM]ACTACTACTACTACTACTACTACTACTACT

Knob-2 (180 bp Knob repeat): [FAM]GAAGGCTAACACCTACGGATTTTTGACCAAGAAATGGTCTCCACCAGAAATCCAAAA AT

MR68-3 (Chromosome 6 clone MR68, AF020265.1): [FAM]CTCTCGTGCGAATACAATGCCCTCAATATCATAGAAACACTATTCCTTTAGTGTGAAT A

CentC69-1 (Centromeric repeat clone CentC69, AY530284.1):

[CY5]CCCAATCCACTACTTTAGGTCCAAAACTCATGTTTGGGGTGGTTTCGCGCAATTTCGT T

Images were taken using an Olympus BX61 microscope equipped with an ORCA-ER CCD camera (Hamamatsu). All images were acquired in grey scale and pseudocoloured with Adobe Photoshop CS5 (Adobe Systems). Karyotyping was done according to [76] using the maize line B73 as a reference. Chromosomes were identified based on distribution of repetitive sequences, morphology and centromeric index.

### Repeat analysis

To obtain equivalent transposon and tandem repeat data for comparative analyses we annotated all six lines with the same annotation workflows. A homology search against the Panicoidae section of the PGSB transposon library [77] resulted in a basal transposon annotation. The used REdat_9.8_Panicoideae contains publicly available Panicoideae transposons templates as well as *de novo* detected full length LTR-retrotranspons from Maize (12,510 elements from B73v2) and Sorghum (3.368 elements). The program vmatch (http://www.vmatch.de), was used as a fast and efficient matching tool well suited for large and highly repetitive genomes under the following parameter setup: identity>=70%, minimal hit length 75 bp, seedlength 12 bp (exact commandline: -d -p -l 75 -identity 70 -seedlength 12 -exdrop 5). The vmatch output was filtered for redundant hits via a priority based approach, which assigns higher scoring matches first and either shortens (<90% coverage and >=50bp rest length) or removes lower scoring overlaps. The resulting annotation is free of overlaps. Elements that have been interrupted by other transposon insertions (nesting) are not defragmented into a higher order instance (like exons belonging to one gene).

Full-length LTR retrotransposons (fl-LTR) where identified with LTRharvest [78] using the following parameters: “overlaps best -seed 30 -minlenltr 100 -maxlenltr 2000 -mindistltr 3000 -maxdistltr 25000 -similar 85 -mintsd 4 -maxtsd 20 -motif tgca -motifmis 1 -vic 60 -xdrop 5 -mat 2 -mis -2 -ins -3 -del -3”. All candidates from the LTRharvest output were subsequently annotated for PfamA domains using hmmer3 (http://hmmer.org) and stringently filtered for false positives by several criteria, the main ones being the presence of at least one typical retrotransposon domain [e.g. reverse transcriptase (RT), RNase H (RH), integrase (INT), protease (PR), etc.)] and a tandem repeat content < 25%. The inner domain order served as a criterion for the classification into the Gypsy (RT-RH-INT) or Copia (INT-RT-RH) superfamily abbreviated as RLG and RLC. Elements missing either INT or RT were classified as RLX. The insertion age of each full-length LTR retrotransposon was estimated based on the accumulated divergence between its 5’ and 3’ long terminal repeats and a random mutation rate of 1.3×10-8 [79].

Tandem repeats where identified with the TandemRepeatFinder under default parameters [80] and subjected to an overlap removal as decribed above prioritizing longer and higher scoring elements. Kmer frequencies were calculated with Tallymer [81].

Syntentic fl-LTRs where identified by sequence clustering (vmatch dbcluster, 98% identity and 98% coverage) of TE-junctions which consisted of 2×100 pb sequence signatures spanning the upstream and downstream insertion sites with each 50bp inside and 50 bp outside of the TE element.

### Whole genome alignments

Block alignments of high identity were generated for all pairwise combinations of the six maize lines using the MUMMER v3 suite [82]. Initial alignments were computed using nucmer with a minimal cluster size of 250 bp (-c 250) and a seed size of 20 bp (-l 20). Results were piped through the delta-filter tool selecting the best one-to-one blocks (global option -1) with a minimal size of 500bp (-l 500). Despite this filter step, a large number (~41-81k) of putatively translocated blocks aligned small regions between different chromosomes or unanchored scaffolds. These blocks frequently overlapped ‘regular’ blocks linking same orthologous chromosomes and showed significantly lower sequence identities in comparison to their respective regular blocks with which they overlapped. Hence, they likely represented paralogous alignments triggered by tandem repeats or the absence of the truly orthologous sequences in one of the lines. To derive the final set of single alignment blocks (SAB), we only included candidate inter-chromosomal translocations with a minimal sequence identity of 99% and a maximum overlap of 10bp to its adjacent alignments. To gain an overview of the contiguity between the six maize genomes, SABs were connected to merged alignment blocks (MAB) if SABs were directly adjacent in both genomes and had a consistent orientation (Supplementary Figure 3). MABs therefore comprise genomic regions of conserved global order and skip genomic parts that were not aligned due to larger sequence differences or undefined sequences (stretches of ‘N’s).

To identify and delineate genomic regions for each line that align (i) to none of the other lines (designated as ‘unaligned’); (ii) to all lines of the same group (either flint or dent maize) and to none of the other group (aka ‘group-specific); (iii) to all five other lines (‘core6’); we superimposed SABs and MABs of one line on its genome coordinates by recording from which group and how many times a single genomic base participated in an alignment. Thereby, we classified each base and concatenated adjacent bases of the same type (i-iii, see above) to derive the unaligned, group-specific and core genomic part. Reversing this approach, identified core regions of EP1 were re-projected onto the pairwise alignments to decode the sequence and coordinates of matching core elements in the other five lines. Based on these coordinates, sequences for each core block were extracted and aligned applying the Fast Sequence Aligner [83] with default parameters. SNPs were directly determined from these multiple sequence alignments (omitting insertions/deletions) and their position in each of the six genomes was derived from the position in the alignment and the known start and end genomic positions for the core block. To deduce higher order haplotypes, we used seeds of the 15 pairwise runs-of-identities that showed for 40 consecutive SNPs an identical haplotype for a particular combination of lines. An iterative greedy algorithm linking the seed with adjacent up- and downstream runs extended such seeds if the run exhibited a haplotype identical to the seed and less than 3 non-matching SNPs were observed between the current end of the extension and the candidate run.

### RNA Preparation and Expression analysis

Different tissues of the European maize (*Zea mays*) inbred lines EP1 and F7 were sampled according to [84, 85] and subsequently subjected to transcriptome sequencing. Supplementary Table 3 provides a detailed description of the collected tissues. Samples 1-3 were collected from seeds imbibed (whole seed, sample 1) or germinated (primary root, sample 2; coleoptile, sample 3) in paper rolls [86] in a 16 h light (28 °C) and 8 h dark (21 °C) regime of a growth cabinet (Conviron CMP6010, http://www.conviron.com). Tissue samples 4-24 were taken from plants grown in a climate chamber either in small pots (14 cm top diameter, 8 cm height, 0,25 l volume, samples 4-17) or in big pots (28 cm top diameter, 21 cm height, 10 l volume, samples 18-24) containing soil substrate type ED 73 (https://www.meyer-shop.com). Growth conditions in the climate chamber were 16 h light (28 °C) and 8 h dark (21 °C). Two individual plants were collected for each of the 24 tissues. Harvested plant material was immediately frozen in liquid nitrogen and stored at −80 °C until RNA extraction. For total RNA extraction, the 24 samples per genotype were separately ground in liquid nitrogen. Subsequently, one spatula of each of the 24 samples per genotype was mixed to generate one pool per inbred line EP1 and F7, respectively. Total RNA was extracted from each of the two pools using the Qiagen RNeasy mini kit according to the manufacturer‘s protocol (Qiagen, https://www.qiagen.com), including on-column DNA digestion. RNA quality was determined by agarose gel electrophoresis and by a Bioanalyzer using an Agilent RNA 6000 Nano Chip (Agilent Technologies, https://www.agilent.com). Both RNA samples were of excellent quality with RNA integrity number values [87] between 9.8 and 10. In total, 9 µg total RNA per pool were used for RNA-seq at Novogene (Novogene Co. Ltd, Bejing, China). After mRNA enrichment using oligo(dT) beads, sequencing libraries were constructed with the NEBNext® Ultra™RNA Library Prep Kit for Illumina® (New England Biolabs, Ipswich, USA), and Illumina PE (150 bp x 2) sequencing was performed on a HiSeq4000 by Novogene. The number of raw reads and quality-filtered reads are shown in Supplementary Table 4. Reads containing adapters, reads containing more than 10% undefined bases and reads containing more than 50% low quality bases (Qscore 5) were removed.

### Expression analysis and analysis of differentially expressed genes (DSG)

We identified regions in SAB, which were fixed within either flint or dent but segregating between the two germplasm of the six lines under study. The corresponding variants were extracted from the Maize Hapmap v3.2.1 panel [38] for lines of the 282 inbred maize association panel [88]. Selected variants were subjected to phylogenetic analysis using FastTree (Version 2.1.5 SSE3) [89] with default parameters and the resulting phylogenetic tree was visualized using iTOL [90] (Supplementary Figure 13). Two non-overlapping subtrees were selected such that three of the lines presented in this study (B73, EP1 and F7) are contained together with other topologically close lines. All lines in the subtree containing B73 were assigned the group one, all those in the subtree containing both EP1 and F7 were assigned group two.

For the resulting subset of of 41 lines (Supplementary Figure 13) samples from seven different tissues (germinating shoot and root, the tip and base of leaves as well as leaf samples under light and dark conditions and kernels) were downloaded from SRA (PRJNA383416) [91]. Trimming of the first thirteen basepairs and cropping of bases with quality values below 20 at the 3’end of reads was performed using bbbduk from the bbmap toolkit [92] as described in the QuantSeq user guide from Lexogen (http://www.bluebee.com/wp-content/uploads/2017/08/QuantSeq-Data-Analysis-Pipeline-User-Guide.pdf).

FastQC (www.bioinformatics.babraham.ac.uk/projects/fastqc) was used before and after trimming to check the quality of the reads. Trimmed reads were mapped to the maize reference sequences B73_v4 using the splice aware STAR aligner (version 2.5.1a) [93]. A read was allowed to map in at most 10 locations (–outFilterMultimapNmax 10) with a maximum of 4% mismatches (–outFilterMismatchNoverLmax 0.04) and all non-canonical intron motifs were filtered out (– outFilterIntronMotifs RemoveNoncanonicalUnannotated). The SAM-format files were converted to BAM-format files and sorted using SAMtools [94]. To obtain non-unique gene-level counts from the mapping files, HTSeq (version 0.9.1) with the ‘nonunique all’-method was used [95]. Normalization of read counts was performed by library sequence depth using the R-package DESeq2 (version 1.23.3) [96]. For differential gene expression analysis with DESeq2 lines classified into either the flint or dent like group were treated as biological replicates. A gene was considered as differentially expressed when the Benjamini-Hochberg adjusted p-value was equal or below 0.05.

### Data availability

The *de novo* assembled genomes and raw reads were released in collaboration with the Maize Genetics and Genomics Database MaizeGDB [18] and NCBI BioProjects PRJNA360923 (F7, DK105 and PE0075) and PRJNA360920 (EP1). Additionally, RNAseq raw data were also deposited under these NCBI accessions.

