## Supplementary material for "European maize genomes unveil pan-genomic dynamics of repeats and genes": european_maize

### Supplementary Tables

**Supplementary Table 1. Read sequences for the assemblies.** Table lists library types, insert sizes, read types, number of libraries and raw read coverage for lines EP1, F7, DK105 and PE0075 used for de novo genome assembly.

| Library type | Insert size | Reads | No. libraries | Coverage EP1 | Coverage F7 | Coverage DK105 | Coverage PE0075 |
|---|---|---|---|---|---|---|---|
| PCR-free PE library (PE250X2) | 450-470 bp | 250 bp x 2 | 1 | 109x | 75x | 68x | 68x |
| PCR-free PE library (PE150X2) | 700-800 bp | 150 bp x 2 | 1 | 84x | 44x | 39x | 39x |
| MP (Nextera™ MP Gel Plus) | 2-4 kbp | 150 bp x 2 | 1 | 34x | 34x | 37x | 41x |
| MP (Nextera™ MP Gel Plus) | 5-7 kbp | 150 bp x 2 | 1 | 34x | 43x | 38x | 33x |
| MP (Nextera™ MP Gel Plus) | 8-10 kbp | 150 bp x 2 | 1 | 59x | 29x | 32x | 36x |
| **Total coverage** | | | | **320x** | **225x** | **214x** | **217x** |

**Supplementary Table 2. Assembly and BUSCO statistics for the four *de novo* assemblies of EP1, F7, PE0075 and DK105.** [1]BUSCO results refer to the Liliopsida dataset.

| | EP1 | F7 | PE0075 | DK105 |
|---|---|---|---|---|
| **DeNovoMAGIC** | *v2* | *v2* | *v3.0* | *v3.0* |
| **Scaffolds stats** | | | | |
| Total scaffolds [#] | 71,196 | 77,899 | 1,288 | 1,393 |
| Assembly size [bp] | 2,462,913,883 | 2,404,712,832 | 2,198,402,809 | 2,288,116,732 |
| Gaps size [bp] | 25,361,986 | 25,595,801 | 14,615,046 | 14,285,020 |
| Gaps [%] | 1.03 | 1.06 | 0.66 | 0.62 |
| N50 [bp] | 6,134,294 | 9,483,449 | 8,642,309 | 10,390,014 |
| MAX [bp] | 29,676,303 | 43,780,026 | 42,352,866 | 40,445,661 |
| **Contigs stats** | | | | |
| Total contigs [#] | 137,249 | 130,426 | 116,681 | 51,270 |
| Assembly size [bp] | 2,434,778,235 | 2,377,026,979 | 2,256,590,598 | 2,242,258,170 |
| N50 [bp] | 82,295 | 96,432 | 109,087 | 101,213 |
| MAX [bp] | 766,959 | 704,566 | 1,314,119 | 1,173,809 |
| **BUSCO[1]** | | | | |
| Complete | 3140 (95.8%) | 3121 (95.2%) | 3139 (95.8%) | 3135 (95.7%) |
| Complete Single-copy | 2761 (84.2%) | 2,718 (82.9%) | 2745 (83.8%) | 2743 (83.7%) |
| Complete Duplicated | 379 (11.6%) | 403 (12.3%) | 393 (12.0%) | 392 (12.0%) |
| Fragmented | 59 (1.8%) | 68 (2.07%) | 57 (1.7%) | 65 (2.0%) |
| Missing | 79 (2.4%) | 89 (2.72%) | 82 (2.5%) | 78 (2.3%) |
| **Total searched** | **3,278** | **3,278** | **3,278** | **3,278** |

**Supplementary Table 3. Overview of sampling for RNAseq** with developmental time points, tissues/organs, number of plants sampled, growth conditions and sampling dates. DAS: days after sowing, DAP: days after pollination. Growth conditions: A – paper rolls, B - soil, small pots, C - soil, big pots.

| Sample no. | Dev. time point | Tissue/organ | No. of plants | Growth cond.* |
|---|---|---|---|---|
| **1** | 24 DAS | germinating whole seed | 2 | A |
| **2** | 4 DAS | primary root | 2 | A |
| **3** | 4 DAS | coleoptile | 2 | A |
| **4** | 9 DAS | seminal & lateral roots | 2 | B |
| **5** | 9 DAS | primary & lateral roots | 2 | B |
| **6** | 9 DAS | pooled leaves | 2 | B |
| **7** | 5-leaf-stage | shoot tip | 2 | B |
| **8** | 5-leaf-stage | topmost leaf | 2 | B |
| **9** | 5-leaf-stage | base of 4th leaf | 2 | B |
| **10** | 5-leaf-stage | tip of 4th leaf | 2 | B |
| **11** | 3-leaf-stage | stem and SAM | 2 | B |
| **12** | 3-leaf-stage | first leaf and sheath | 2 | B |
| **13** | 3-leaf-stage | topmost leaf | 2 | B |
| **14** | 3-leaf-stage | all roots (primary, lateral, seminal roots) | 2 | B |
| **15** | 5-leaf-stage | crown roots | 2 | B |
| **16** | 5-leaf-stage | seminal & lateral roots | 2 | B |
| **17** | 5-leaf-stage | primary & lateral roots | 2 | B |
| **18** | 1 day before pollination (R1) | non-flowering tassel | 2 | C |
| **19** | 1 day before pollination (R1) | silk | 2 | C |
| **20** | 1 day before pollination (R1) | innermost husk | 2 | C |
| **21** | 1 day before pollination (R1) | uppermost leaf | 2 | C |
| **22** | 1 day before pollination (R1) | shoot-borne roots | 2 | C |
| **23** | 1 day before pollination (R1) | pre-pollination comb | 2 | C |
| **24** | 4 DAP | whole seed | 2 | C |

**Supplementary Table 4. RNAseq quality summary.** Raw and clean bases are denoted by Gb.

| Sample | Raw Reads | Clean Reads | Raw bases | Clean bases | Effective rate (%) | Error rate (%) | Q20 (%) | Q30 (%) | GC content (%) |
|---|---|---|---|---|---|---|---|---|---|
| **F7** | 688499867 | 667448978 | 206.55 | 200.23 | 96.94 | 0.02 | 95.31 | 89.19 | 56.60 |
| **EP1** | 692842987 | 652393710 | 207.85 | 195.72 | 94.16 | 0.01 | 95.92 | 90.25 | 56.45 |

**Supplementary Table 5. Sequence accuracy for EP1 and F7 assemblies.** Illumina reads of 30 maize lines (NCBI Bioproject PRJNA260788) were mapped by BWA to the respective reference genomes and SNPs were called using bcftools with no filtering thereby reporting all possibly variant sites. To avoid mapping artifacts for the evaluation of the EP1 and F7 sequence accuracy, we scored only sites with a read depth ≥ 10, mapping quality ≥ 20, genotype quality GQ ≥ 10 and no strand bias (SBphred == 0). Columns show per chromosome numbers of total scored sites (‘sites’), and calls that show no (‘consistent’), a homozygous (‘homoalt’) or heterozygous (‘hetalt’) difference to the reference sequence, respectively.

| | EP1 | | | | F7 | | | |
|---|---|---|---|---|---|---|---|---|
| scaffold | sites | consistent | homoalt | hetalt | sites | consistent | homoalt | hetalt |
| chr_1 | 7589219 | 7589158 | 8 | 53 | 8129700 | 8129664 | 9 | 27 |
| chr_2 | 6037276 | 6037246 | 4 | 26 | 6309689 | 6309658 | 6 | 25 |
| chr_3 | 5920655 | 5920620 | 1 | 34 | 6256886 | 6256850 | 4 | 32 |
| chr_4 | 6671784 | 6671707 | 17 | 60 | 6853201 | 6853156 | 13 | 32 |
| chr_5 | 5396328 | 5396296 | 3 | 29 | 5602100 | 5602074 | 2 | 24 |
| chr_6 | 4002220 | 4002194 | 2 | 24 | 4209873 | 4209853 | 1 | 19 |
| chr_7 | 4502468 | 4502434 | 12 | 22 | 4778508 | 4778484 | 1 | 23 |
| chr_8 | 4427095 | 4427072 | 1 | 22 | 4653156 | 4653133 | 2 | 21 |
| chr_9 | 4144162 | 4144116 | 4 | 42 | 4287042 | 4287010 | 11 | 21 |
| chr_10 | 3825773 | 3825746 | 1 | 26 | 3942482 | 3942455 | 1 | 26 |
| **ALL** | **52516980** | **52516589** | **53** | **338** | **55022637** | **55022337** | **50** | **250** |

**Supplementary Table 6. Repeat composition of six maize lines. A)** Transposons detected via homology to REdat_9.8_Panicoideae, without overlapping annotations and as percent of the respective assembly length without Ns. Overall and subgroup numbers are very similar between all lines. Even PH207 is not much different in its transposon content, despite its lower assembly assembly quality. **B)** Simple sequence tandem repeats and subgroups in Mb (overlaps removed). The large up to 10 fold differences in satellite and knob tandem repeats reflect different assembly strategies and not biological differences as show by a fish analyses.

A

| (% of Nfree assembly) | B73v4 | PH207 | EP1 | F7 | DK105 | PE0075 |
|---|---|---|---|---|---|---|
| **Mobile Element (TXX)** | **79.5** | **77.2** | **81.1** | **80.7** | **80.4** | **80.0** |
| **Class I: Retroelement (RXX)** | **77.2** | **74.9** | **79.1** | **78.6** | **78.3** | **77.8** |
| LTR Retrotransposon (RLX) | 76.8 | 74.5 | 78.8 | 78.3 | 77.9 | 77.5 |
| Ty1/copia (RLC) | 22.4 | 18.4 | 24.6 | 24.0 | 22.8 | 22.8 |
| Ty3/gypsy (RLG) | 37.4 | 36.3 | 36.7 | 36.7 | 37.6 | 37.4 |
| unclassified LTR (RLX) | 17.0 | 19.8 | 17.5 | 17.6 | 17.5 | 17.3 |
| non-LTR Retrotransposon (RXX) | 0.37 | 0.44 | 0.32 | 0.34 | 0.35 | 0.36 |
| LINE (RIX) | 0.37 | 0.44 | 0.32 | 0.34 | 0.35 | 0.36 |
| **Class II: DNA Transposon (DXX)** | **2.28** | **2.29** | **2.03** | **2.07** | **2.11** | **2.16** |
| DNA Transposon Superfamily (DTX) | 1.34 | 1.19 | 1.20 | 1.21 | 1.23 | 1.26 |
| CACTA superfamily (DTC) | 1.00 | 0.81 | 0.92 | 0.91 | 0.92 | 0.93 |
| hAT superfamily (DTA) | 0.10 | 0.11 | 0.09 | 0.09 | 0.09 | 0.09 |
| Mutator superfamily (DTM) | 0.12 | 0.11 | 0.10 | 0.11 | 0.11 | 0.11 |
| Tc1/Mariner superfamily (DTT) | 0.001 | 0.001 | 0.001 | 0.001 | 0.001 | 0.001 |
| PIF/Harbinger (DTH) | 0.11 | 0.14 | 0.09 | 0.10 | 0.10 | 0.11 |
| unclassified (DTX) | 0.01 | 0.01 | 0.01 | 0.01 | 0.01 | 0.01 |
| MITE (DXX) | 0.55 | 0.68 | 0.48 | 0.49 | 0.51 | 0.53 |
| Helitron (DHH) | 0.18 | 0.23 | 0.16 | 0.15 | 0.17 | 0.17 |
| unclassified DNA transposon (DXX) | 0.20 | 0.20 | 0.19 | 0.21 | 0.20 | 0.20 |
| *Retro-TE/DNA-TE ratio* | *33.9* | *32.7* | *38.9* | *38.1* | *37.2* | *36.0* |
| *Gypsy/Copia ratio* | *1.7* | *2.0* | *1.5* | *1.5* | *1.6* | *1.6* |

B

| (Mb) | B73v4 | PH207 | EP1 | F7 | DK105 | PE0075 |
|---|---|---|---|---|---|---|
| Tandem repeats | 67.4 | 50.5 | 121.2 | 109.0 | 93.8 | 81.7 |
| Microsatellite (2-9bp units) | 1.5 | 0.8 | 1.6 | 1.5 | 1.3 | 1.4 |
| Minisatellite (10-99 units) | 49.7 | 34.3 | 57.6 | 56.2 | 52.8 | 51.3 |
| Satellite (>=100 bp units) | 16.3 | 15.4 | 62.0 | 51.3 | 39.7 | 28.9 |
| knob | 2.62 | 4.06 | 29.83 | 27.29 | 14.40 | 9.61 |

**Supplementary Table 7. Characteristics of *de novo* detected full length LTR retrotransposons in six maize lines. A)** Main detection metrics. **B)** Percent of shared syntenic locations for still intact full length elements, per superfamily and overall. Almost half of all fl-LTR locations are unique to one line. These structural differences are caused by very recent line specific insertions as well as by line specific removals or truncations and may additionally be biased by differing assembly approaches.

A

| | candidates | quality filtered | % of candidates | RLC | RLG | RLX |
|---|---|---|---|---|---|---|
| B73v4 | 78,275 | 14,777 | 18.9 | 3,020 | 5,068 | 6,689 |
| PH207 | 59,529 | 6,838 | 11.5 | 1,141 | 2,162 | 3,535 |
| EP1 | 85,072 | 14,946 | 17.6 | 2,862 | 4,697 | 7,387 |
| F7 | 83,983 | 14,535 | 17.3 | 2,698 | 4,656 | 7,181 |
| DK105 | 81,327 | 14,489 | 17.8 | 2,832 | 4,788 | 6,869 |
| PE0075 | 80,501 | 14,827 | 18.4 | 2,873 | 4,970 | 6,984 |

B

| | % of all fl-LTRs from 6 maize lines | | | |
|---|---|---|---|---|
| **shared between** | **RLC** | **RLG** | **RLX** | **ALL** |
| unique to 1 line | 10.3 | 18.5 | 18.4 | 47.3 |
| 2 lines | 4.3 | 6.4 | 10.3 | 21.0 |
| 3 lines | 2.4 | 3.6 | 7.8 | 13.7 |
| 4 lines | 1.3 | 2.4 | 5.4 | 9.1 |
| 5 lines | 0.8 | 1.2 | 3.9 | 5.9 |
| 6 lines | 0.2 | 0.6 | 2.2 | 3.0 |
| sum | 19.2 | 32.8 | 48.0 | 100 |

**Supplementary Table 8. Total SAB sizes of the pairwise alignments of the six lines.** Rows indicate the target genome line, columns the source genome aligned to the target. For example, the total size of SABs in the PH207 genome that has been detected by B73 (v4) genomic sequences is 1062 Mb. The highest proportion of aligned genomic regions are observed between lines of the same germplasm. Total SAB spans including PH207 are generally decreased due to the significantly larger gap sequences of this line. Note that total sizes can be slightly asymmetric due to differences in alignment gaps.

| | B73v4 | PH207 | EP1 | F7 | DK105 | PE0075 |
|---|---|---|---|---|---|---|
| B73v4 | 0 | 1063 | 1156 | 1182 | 1177 | 1200 |
| PH207 | 1062 | 0 | 970 | 973 | 948 | 1003 |
| EP1 | 1156 | 970 | 0 | 1282 | 1246 | 1225 |
| F7 | 1182 | 973 | 1282 | 0 | 1291 | 1241 |
| DK105 | 1177 | 948 | 1246 | 1291 | 0 | 1275 |
| PE0075 | 1200 | 1003 | 1225 | 1241 | 1275 | 0 |

Total SAB size [Mb]

**Supplementary Table 9. Total sizes of pairwise MABs**, see legend Supplementary Table 7 for details.

| | B73v4 | PH207 | EP1 | F7 | DK105 | PE0075 |
|---|---|---|---|---|---|---|
| B73v4 | 0 | 1838 | 1802 | 1807 | 1798 | 1805 |
| PH207 | 1742 | 0 | 1695 | 1706 | 1676 | 1708 |
| EP1 | 1955 | 1933 | 0 | 2013 | 1998 | 1981 |
| F7 | 1924 | 1905 | 1973 | 0 | 1967 | 1942 |
| DK105 | 1893 | 1865 | 1934 | 1955 | 0 | 1940 |
| PE0075 | 1834 | 1837 | 1860 | 1863 | 1874 | 0 |

Total MAB size [Mb]

**Supplementary Table 10. Number of SNPs and total genomic size of the 31 higher order haplotypes.** Group1 and group2 show the line combinations for which the haplotype is identical within each group and different between the groups. Further descriptions are provided in the main text and methods section.

| Group 1 | Group 2 | SNP count | Size [bp] |
|---|---|---|---|
| B73v4 | DK105;EP1;F7;PE0075;PH207 | 114041 | 31530980 |
| B73v4;DK105;F7;PE0075;PH207 | EP1 | 101909 | 24354517 |
| B73v4;DK105;EP1;F7;PE0075 | PH207 | 90947 | 21225493 |
| B73v4;DK105;EP1;PE0075;PH207 | F7 | 86032 | 23546344 |
| B73v4;DK105;EP1;F7;PH207 | PE0075 | 81117 | 20572261 |
| B73v4;EP1;F7;PE0075;PH207 | DK105 | 71501 | 18687278 |
| **B73v4;PH207** | **DK105;EP1;F7;PE0075** | **71087** | **18763719** |
| B73v4;PE0075;PH207 | DK105;EP1;F7 | 54276 | 14924194 |
| B73v4;F7;PE0075;PH207 | DK105;EP1 | 42024 | 9423938 |
| B73v4;EP1;PE0075;PH207 | DK105;F7 | 38763 | 9487100 |
| B73v4;EP1;PH207 | DK105;F7;PE0075 | 28113 | 6723233 |
| B73v4;DK105;EP1;F7 | PE0075;PH207 | 26926 | 6972824 |
| B73v4;F7;PH207 | DK105;EP1;PE0075 | 26594 | 9456847 |
| B73v4;DK105 | EP1;F7;PE0075;PH207 | 24046 | 6517900 |
| B73v4;DK105;EP1;PE0075 | F7;PH207 | 23052 | 4879837 |
| B73v4;EP1;F7;PH207 | DK105;PE0075 | 22231 | 6282974 |
| B73v4;DK105;PE0075;PH207 | EP1;F7 | 20518 | 5071033 |
| B73v4;DK105;F7;PE0075 | EP1;PH207 | 20513 | 4213886 |
| B73v4;DK105;EP1;PH207 | F7;PE0075 | 19468 | 5462675 |
| B73v4;DK105;F7;PH207 | EP1;PE0075 | 18091 | 4060362 |
| B73v4;DK105;PH207 | EP1;F7;PE0075 | 16329 | 4362025 |
| B73v4;PE0075 | DK105;EP1;F7;PH207 | 16251 | 5553633 |
| B73v4;DK105;EP1 | F7;PE0075;PH207 | 14082 | 2937247 |
| B73v4;EP1;PE0075 | DK105;F7;PH207 | 12244 | 4088402 |
| B73v4;F7 | DK105;EP1;PE0075;PH207 | 12078 | 2811297 |
| B73v4;EP1 | DK105;F7;PE0075;PH207 | 11023 | 2804651 |
| B73v4;DK105;PE0075 | EP1;F7;PH207 | 10456 | 2645995 |
| B73v4;F7;PE0075 | DK105;EP1;PH207 | 10402 | 3669615 |
| B73v4;DK105;F7 | EP1;PE0075;PH207 | 9862 | 2362083 |
| B73v4;EP1;F7;PE0075 | DK105;PH207 | 8412 | 2112371 |
| B73v4;EP1;F7 | DK105;PE0075;PH207 | 8041 | 2639432 |
| | | **1110429** | **288144146** |

### Supplementary Figures

**Supplementary Figure 1. Genetic versus physical distances for all ten chromosomes of the EP1xPH207 cross.** X-axis: physical position on the respective EP1 chromosome in [Mb], Y-axis: genetic position in centiMorgan [cM].

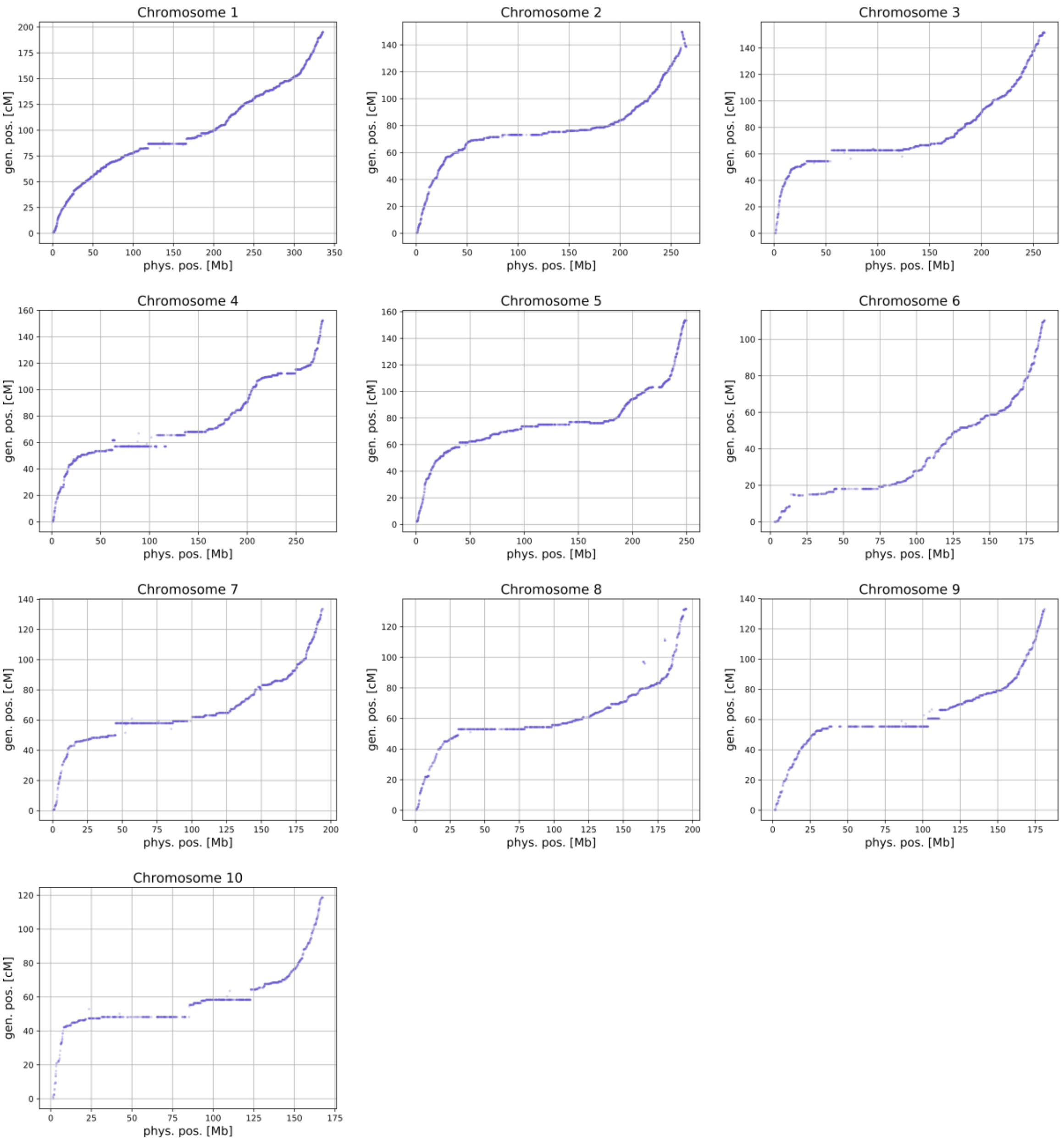

**Supplementary Figure 2. Assembly sizes and overall repeat content of 6 maize lines. A)** Chromosome sizes and and N-content (black bars), **B)** Basic repetitivity in form of 20mer frequencies. Cumulative plot, lower curves represent higher repetitivity e.g. 50% of the PH207 assembly consists of 20mers occurring <=10 times, for the other lines this value is only around 40%. **C)** Transposon content. More details are given in Supplementary Table 6A.

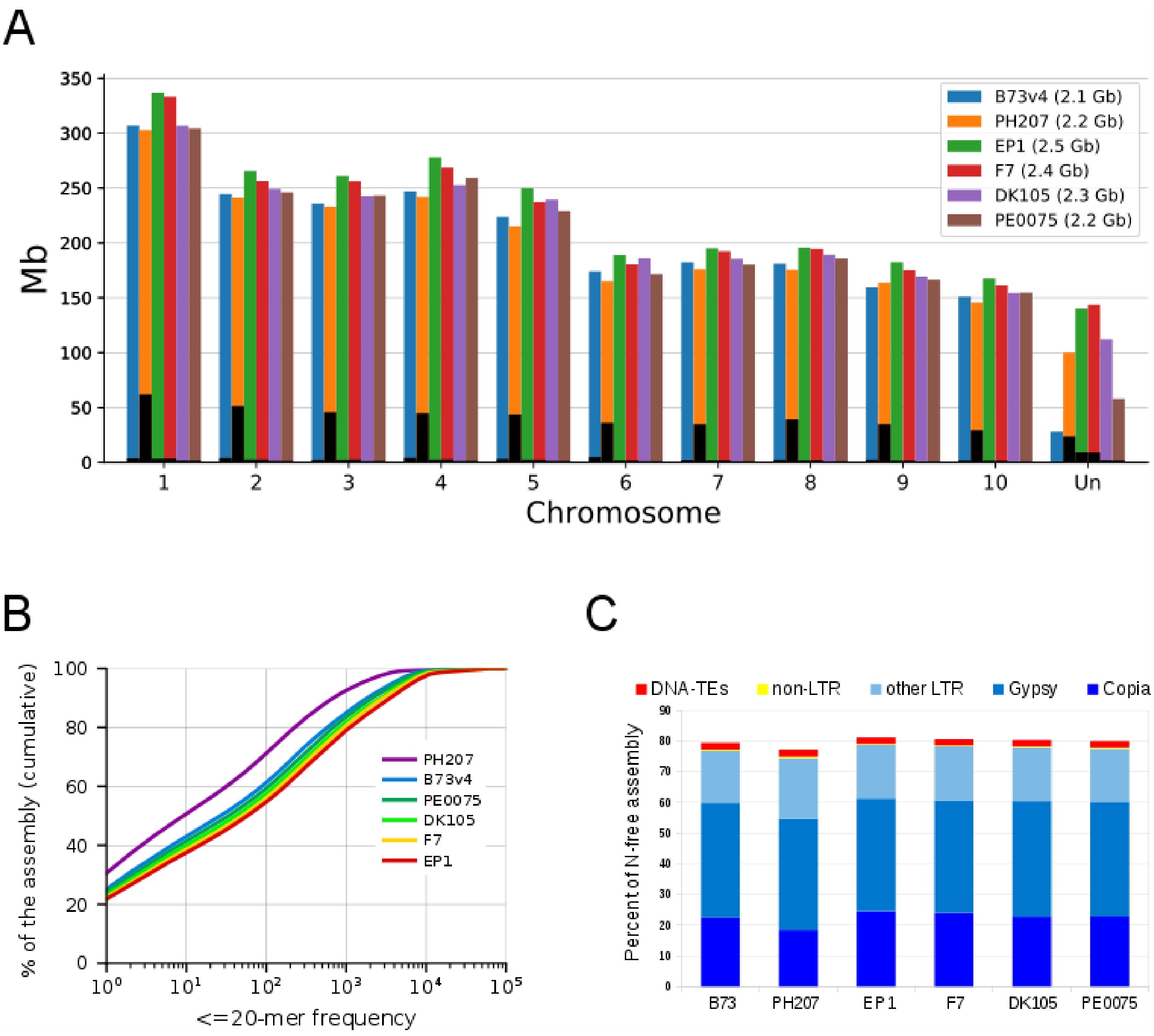

**Supplementary Figure 3**. **Full length LTR-retrotransposon detection in six maize lines. A)** *De novo* detection and stringent quality filtering of fl-LTR elements. **B)** Retrieved fl-LTR numbers as proxy for assembly quality. Five of the six assemblies contain the expected number of fl-LTRs. The by 50% reduced numbers of high quality elements in PH207 reflect an older not optimized assembly approach. **C)** Insertion age distribution of fl-LTRs. The four NRGene assemblies have very similar age distribution patterns. The B73 assembly has a higher proportion of very young Copia and Gypsy elements, which can probably be attributed to technical assembly differences. In PH207, the young elements are completely missing in the assembly and thus explain the reduced numbers. Due to their (almost) identical long terminal repeats the correct structure of young elements is more difficult to resolve.

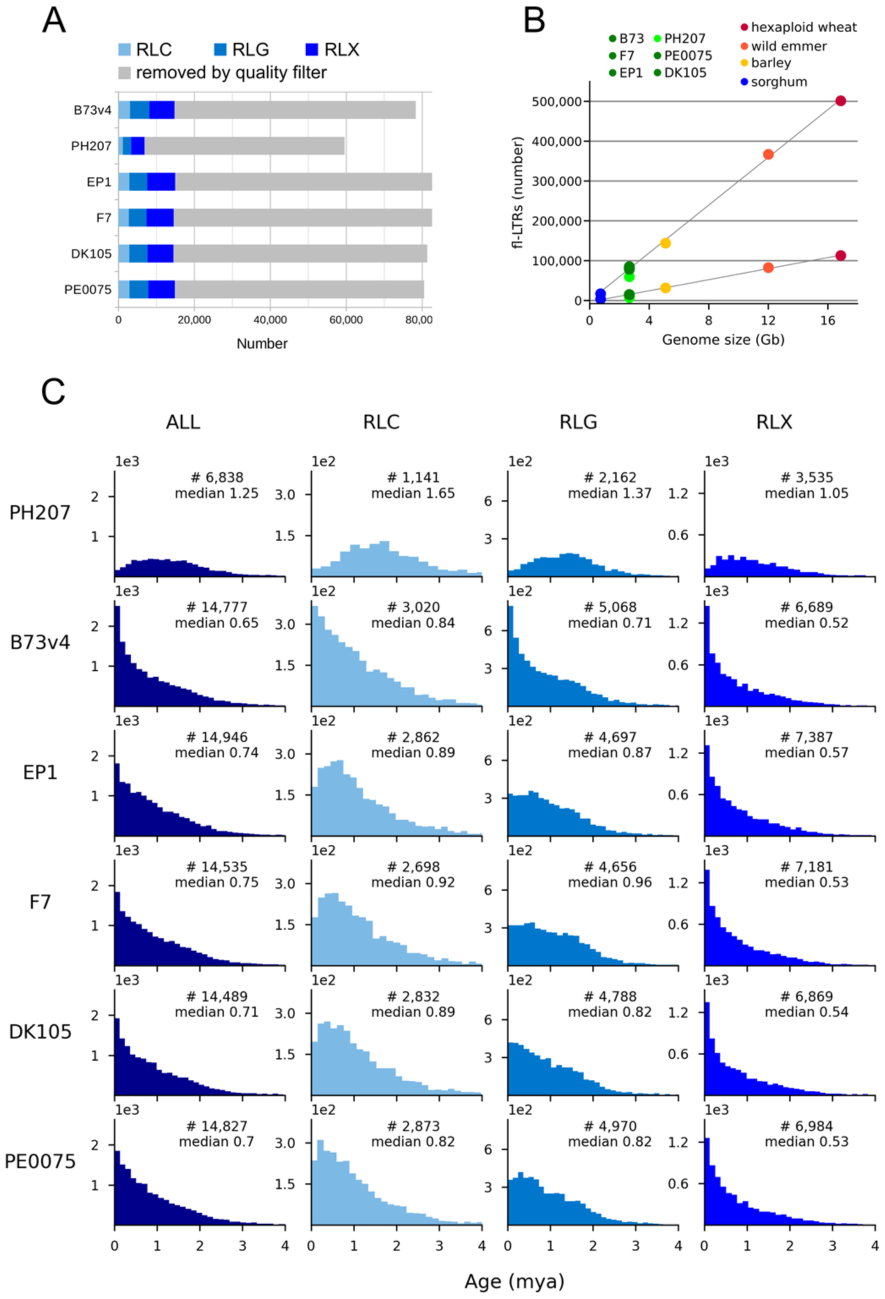

**Supplementary Figure 4. Chromosomal distribution of satellite tandem repeats in the assemblies of six maize lines.** The four NRGene assemblies contain larger amounts of satellite tandem repeats compared to the B73 and PH207 assemblies (Figure 2B). For EP7 they have even been placed in their correct chromosomal context on chromosomes 4L, 5L, and 6S as proven by the fish data (Figure 2A). For DK105 and PE0075 the 5L and 6S locations are present in the assembly, but not the prominent 9S location.  Most of the highly repetitive satellite sequences could not be assigned to a specific chromosomal position. They have been merged into the unassigned sequence pool (Un).

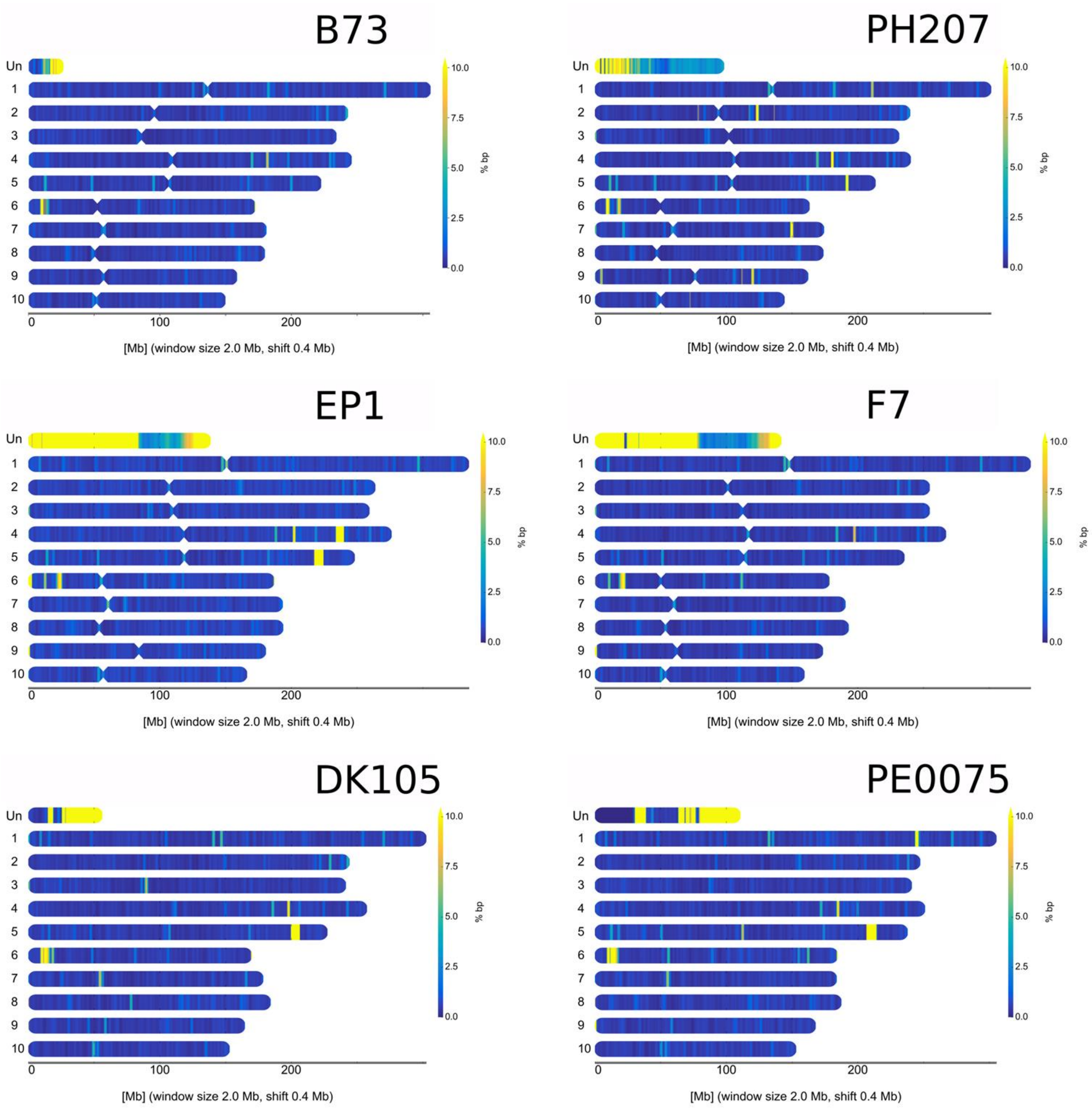

**Supplementary Figure 5. Tandem repeat content and composition in the genome assemblies. A)** The assemblies of EP1 and F7 captured the highest amounts of tandem repeats and knob sequences. The in spite of high fish intensities much lower contents in the B73 and PH207 assemblies reflect the assembly difficulties of these highly repetitive genome regions. **B)** Relationship between the six lines derived from knob location similarities. The input matrix for clustering contained three relative intensities of fish knob locations per chromosome. The knob patterns of the two Dents B73 and PH207 are very similar, as well as the patterns of DK105 and PE0075. **C)** Multiple sequence alignment of selected knob monomers from EP1 together with known knob monomers. The knob sequences in the EP1 assembly consist of 180 and 202 bp monomers with a surplus of the 180 bp monomer by a factor of 6.8. Both monomers are highly similar to previously reported knob monomers from maize with the following Genbank IDs marked as 1, 2, 3 in the figure, 1: AF030934.1, 2: M32521.1 and M32525.1, 3: DQ352544.1_a and DQ352544.1_b. Consensus sequences of the monomers where used to identify all major and minor knob locations in the assemblies. **D)** Gene expression (maximal expression of 7 different conditions per Gene, log10) in relation to the nearest upstream knob signature (left) and satellite tandem repeat (right). Both axes are logarithmic.

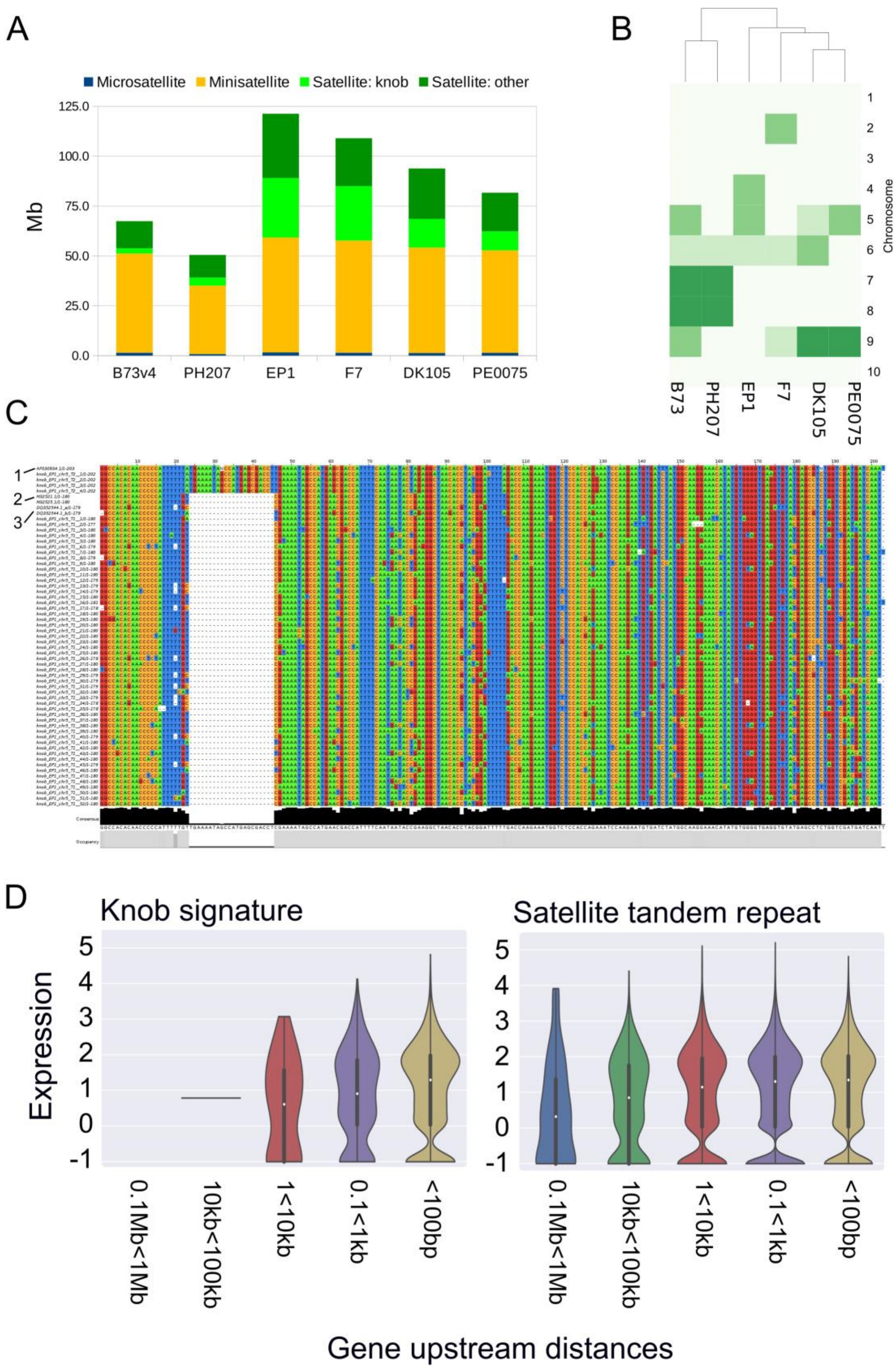
A
Microsatellite
Minisatellite
Satellite: knob
Satellite: other
Mb
125.0
100.0
75.0
50.0
25.0
0.0
B73v4
PH207
EP1
F7
DK105
PE0075
B
Chromosome
1
2
3
4
5
6
7
8
9
10
B73
PH207
EP1
F7
DK105
PE0075
C
1
2
3
Consensus
Occupancy
D
Knob signature
Satellite tandem repeat
Expression
5
4
3
2
1
0
-1
0.1Mb<1Mb
10kb<100kb
1<10kb
0.1<1kb
<100bp
Gene upstream distances

**Supplementary Figure 6. Sketch illustrating the chaining of single alignment blocks (SABs) to merged alignment block (MABs).** SABs were computed as global (1:1) orthologous blocks using the MUMMER tool (see methods). MABs are chains of consistent SABs (upper panel) which can be linked by no edit operation (e.g. insertion, translocation, inversion). The lower panel provides examples of non-permissive edit operations.

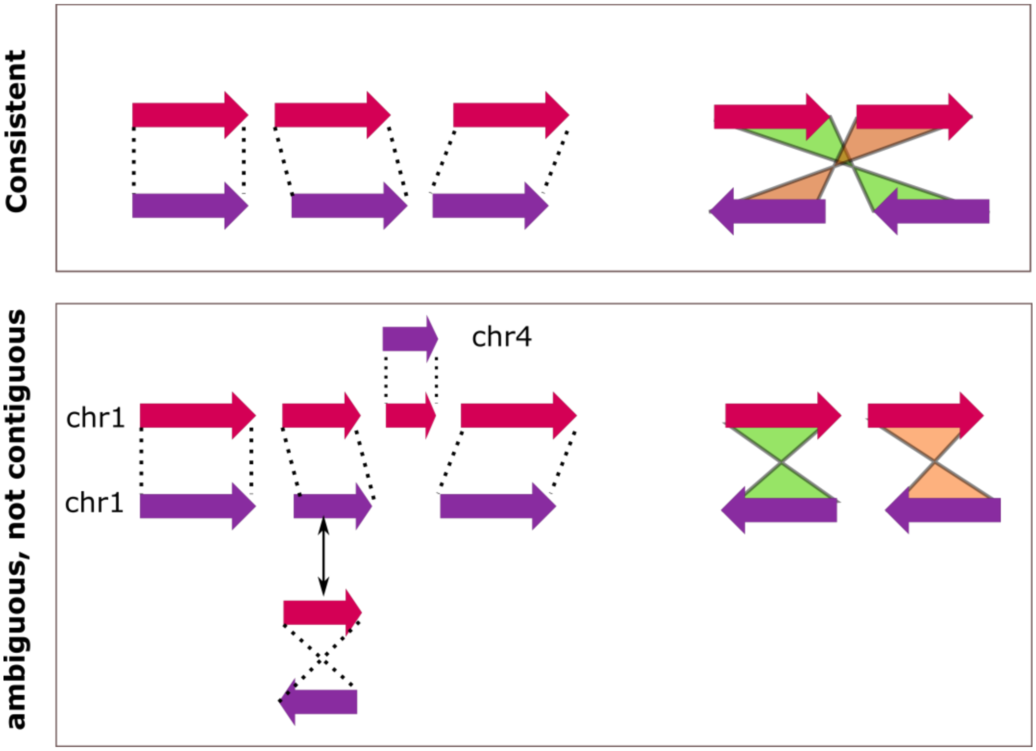

**Supplementary Figure 7. SAB and MAB sizes for all 15 pairwise WGA comparisons** of the six maize lines EP1, F7, DK105, PE0075, B73 (version 4) and PH207. Mean and median are indicated in the boxplot by an asterisk and yellow line, respectively.

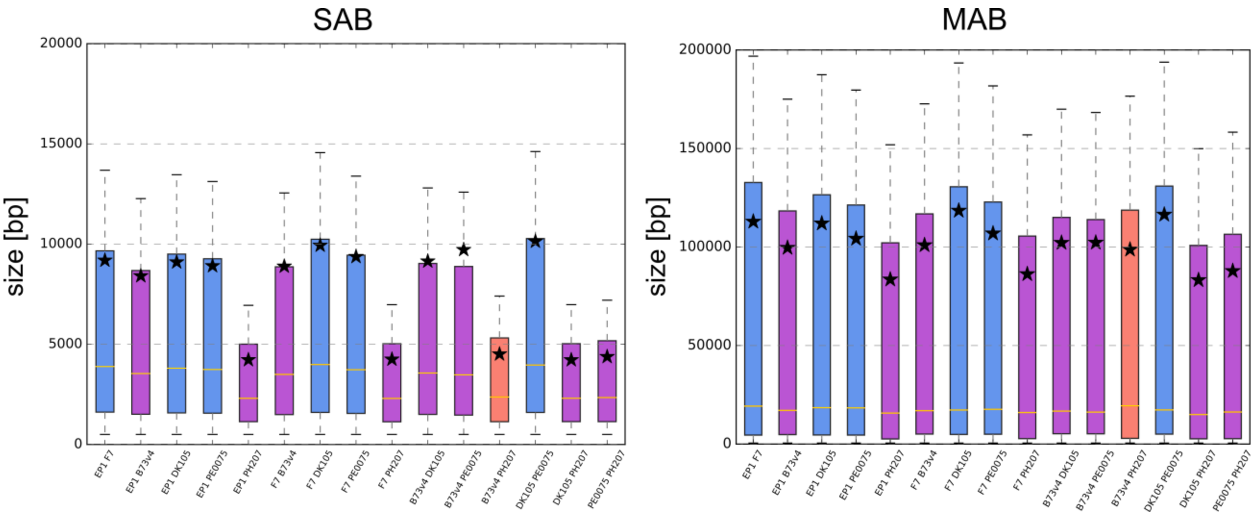

**Supplementary Figure 8. Sizes of inversions (top panel) and translocations (lower panel) for all 15 pairwise WGA comparisons of the six maize lines EP1, F7, DK105, PE0075, B73 (version 4) and PH207.** X-axis: line pair for comparison, Y-axis: size of inversions/translocations in log10 [bp].

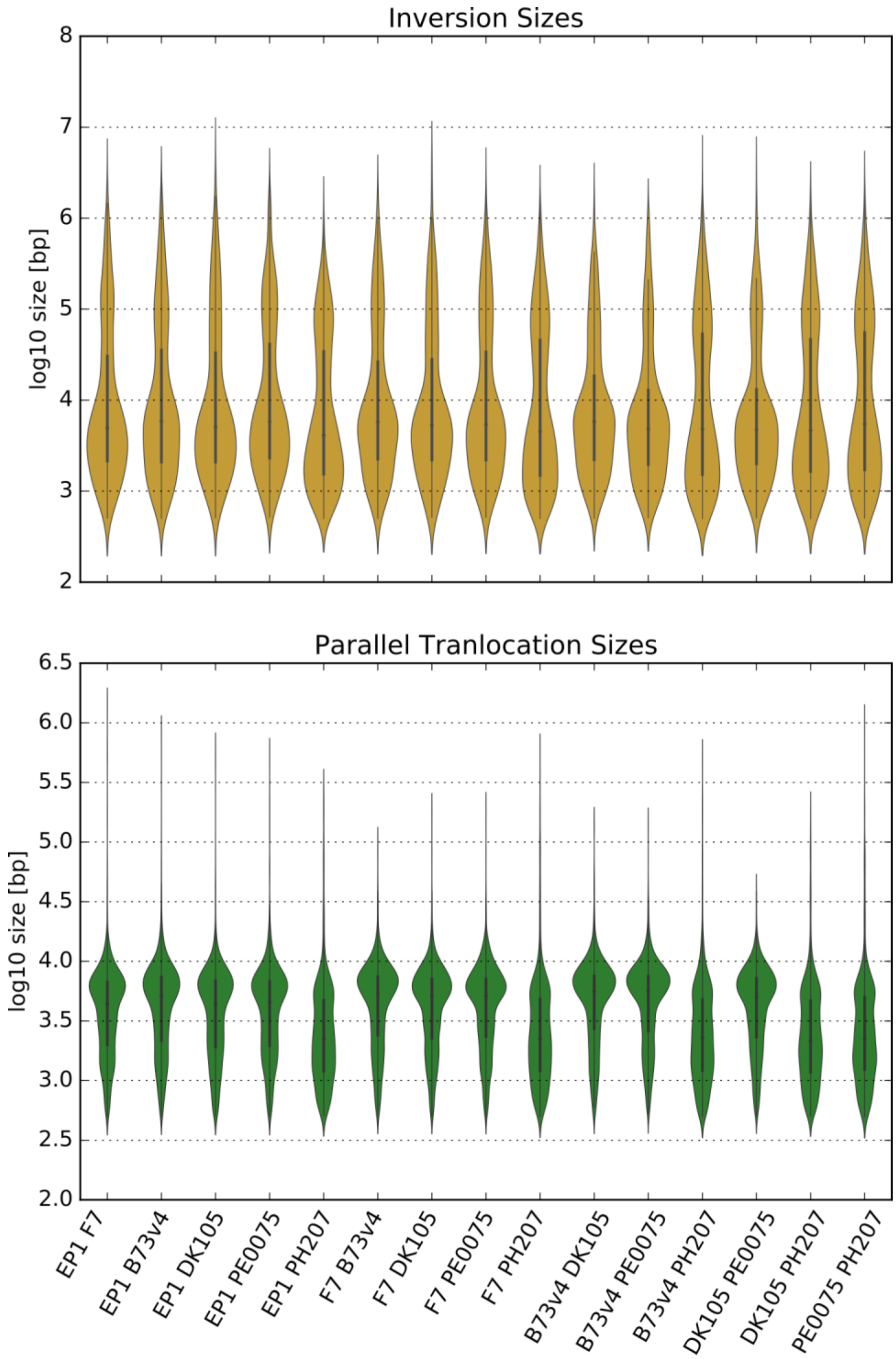

**Supplementary Figure 9. Concept of the genomic core-, group-specific and unaligned regions generated from pairwise WGAs.** Pairwise alignments of EP1 with B73, PH207, F7, DK105 and PE0075 are projected onto the EP1 sequence providing a uniform coordinate system. Such a projection can be performed for MABs (upper panel) and SABs (lower panel). Regions delineated as unaligned regions, group-specific and core regions are labeled 1, 2 and 3,respectively. Note that labeling can substantially vary between MABs or SABs due to the ability of MABs to span unaligned regions between two SABs as long as they can be contiguously linked (see Supplementary Figure 6).

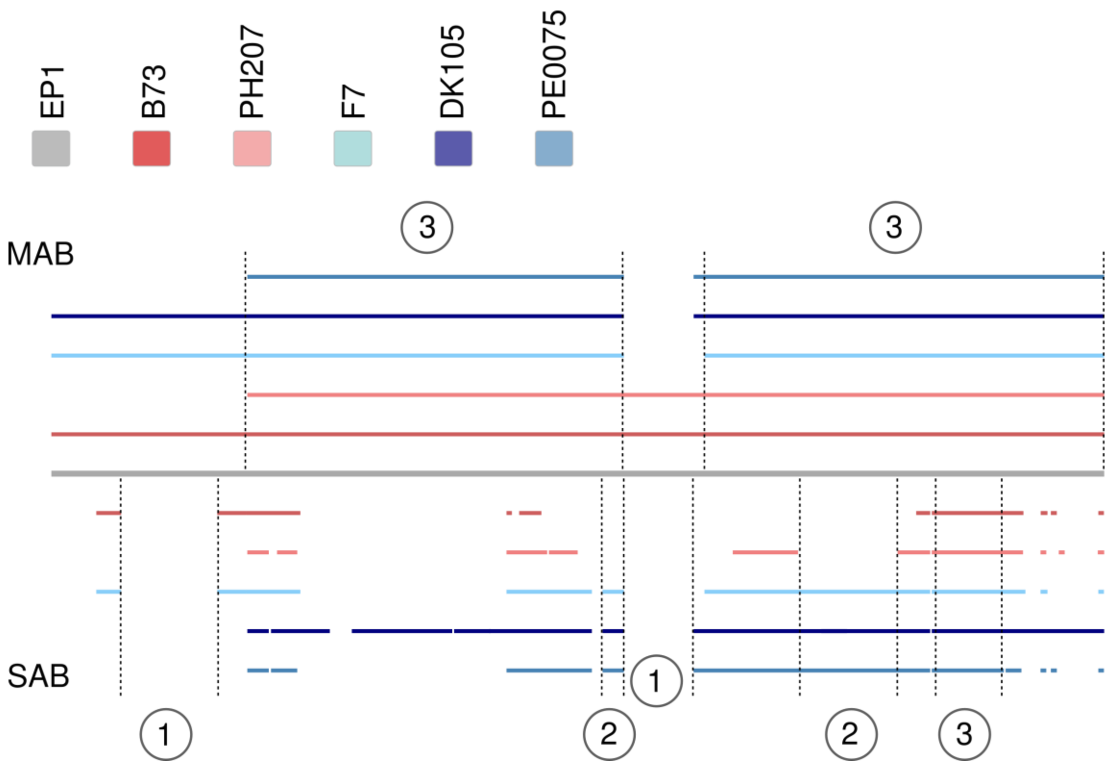

**Supplementary Figure 10. Densities of the core, group-specific and unaligned regions for the six maize lines studies.** Intensities are normalized within one track, dark red/blue values representing a higher fraction of aligned regions per 5 Mb bins. Outer chromosome ideograms show the largest of all six lines, dark grey ellipses indicate the approximate centromere position derived from the repeat analysis. Group of six circular tracks show from outer to inner the core, germplasm-specific and unaligned densities. Each group consists of six tracks, displaying from outer to inner densities for B73, PH207, EP1, F7, DK105 and PE0075.

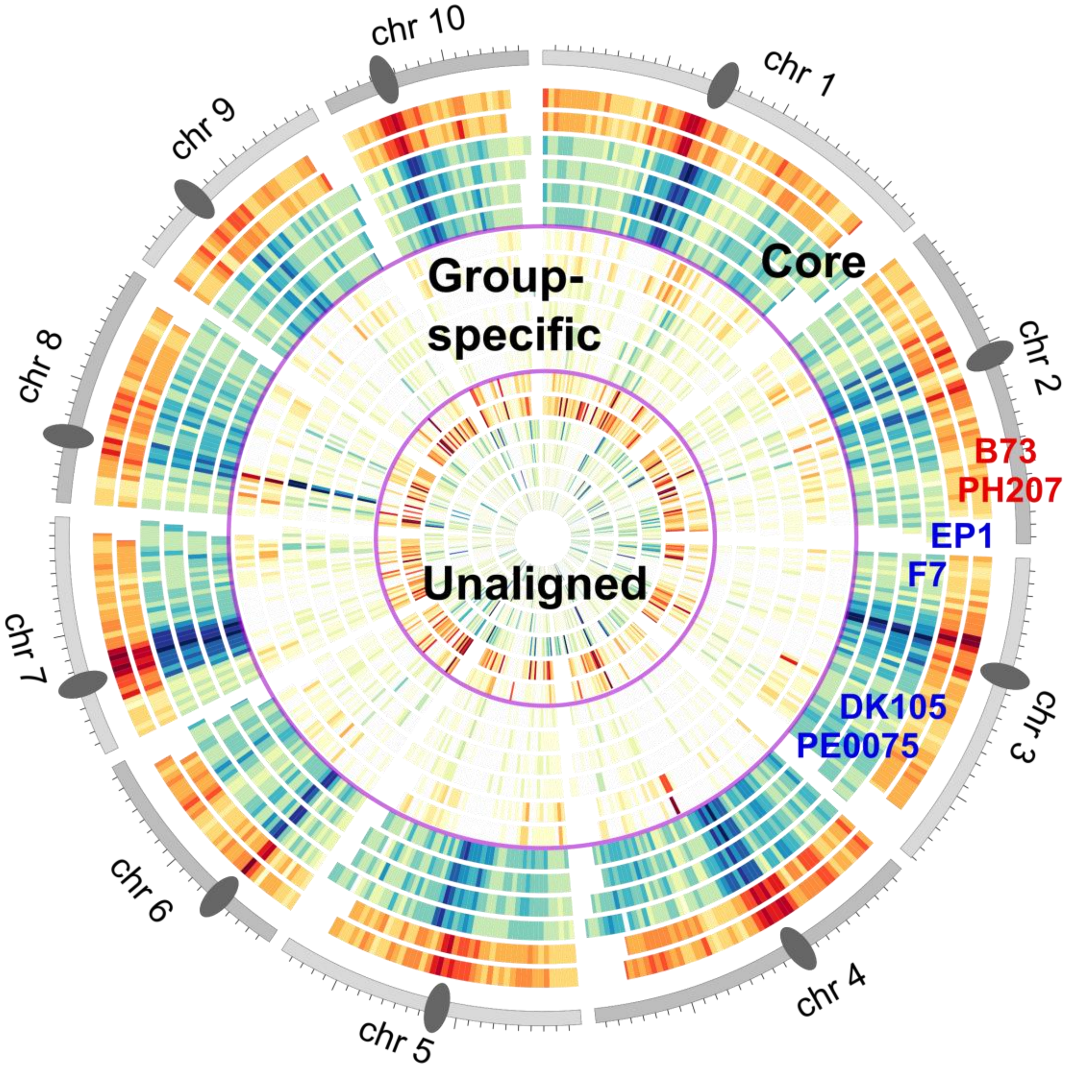

**Supplementary Figure 11. Genomic sizes in [Mb] of SNP runs-of-identity (RoI) for all 15 pairwise comparisons of the six maize lines.** Comparisons within one germplasm (flint: EP1, F7, DK105 and PE0075; dent: B73 version 4 and PH207) share larger genomic fractions than inter-germplasm RoI totals. Y-axis: total sum of RoIs in [Mb], X-axis: line pairs, grey circles indicate maize line used in the respective comparison.

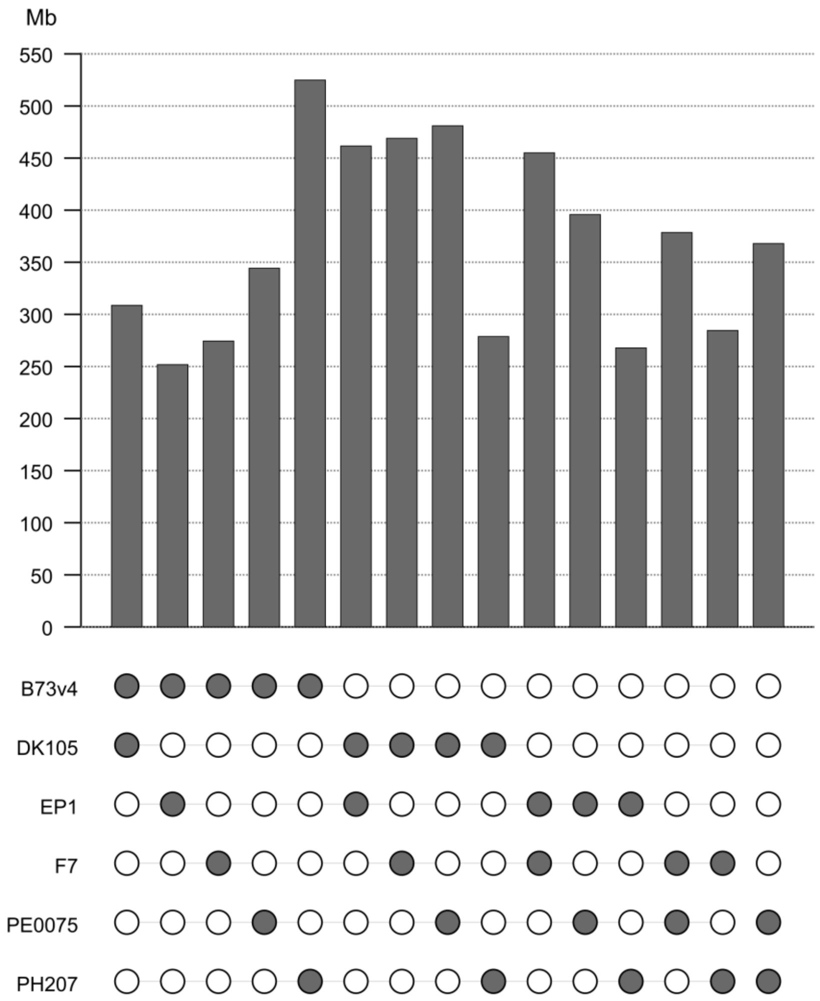

**Supplementary Figure 12. Group-specific syntelogs for all (4,2)-combinations of the six lines and their syntelog set.** We divided the six maize lines of this study into two groups of 4 (gray circles) and 2 (white circles) lines, respectively. For all 15 combinations, the number of group-specific syntelogs for each group was determined under the prior of the identified orthologous gene set. The germplasm specific separation of flint (EP1, F7, DK105 and PE0075) versus dent (B73 version 4 and PH207) genotypes showed the largest distinction between the two groups.

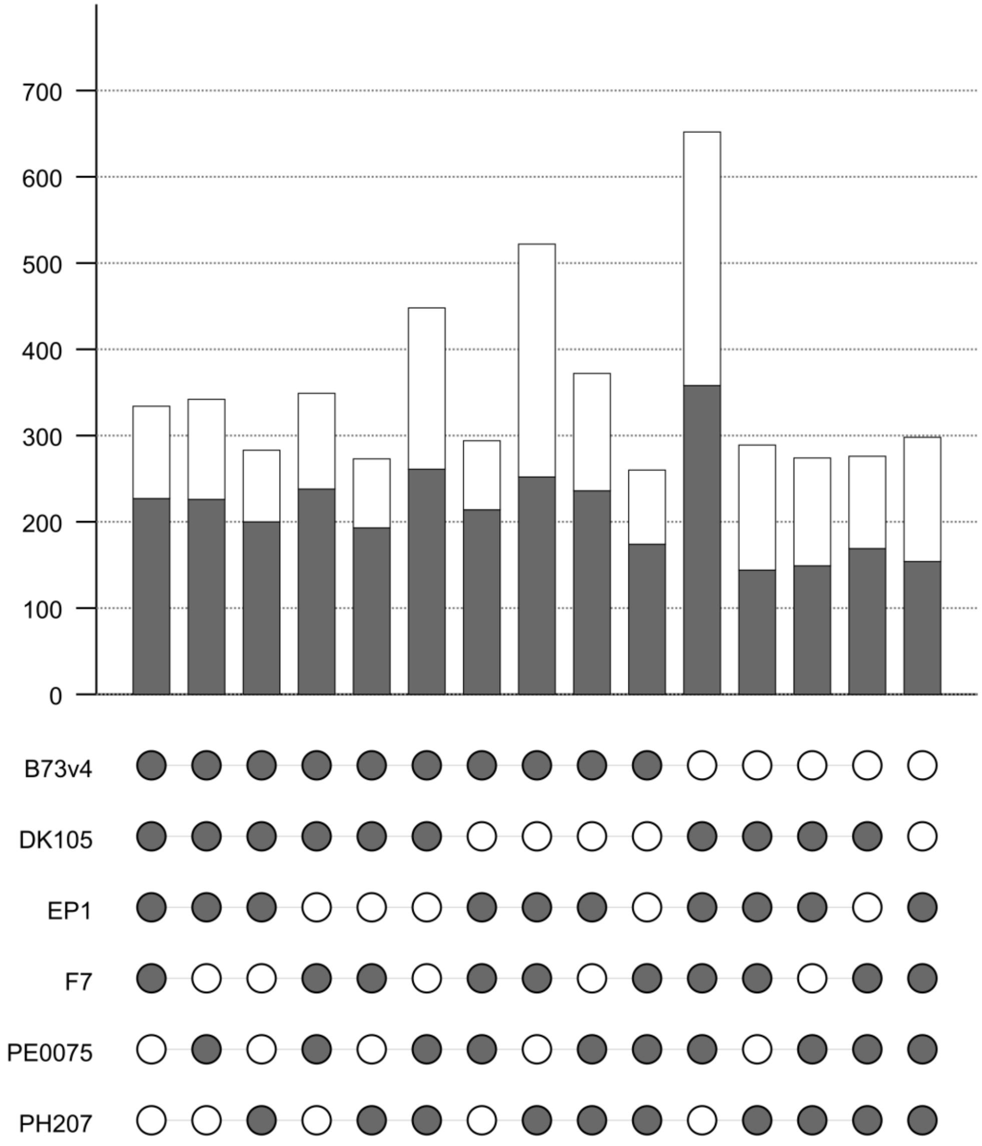

**Supplementary Figure 13. Lines selected for the European flint-dent haplotype expression analysis.** Hapmap v3 SNPs were restricted to the genomic regions defined by the haplotype differentiating European flint lines and dent lines. Based on these variation data, a phylogenetic analysis detected NAM maize lines closely related to B73 (red) or EP1/F7 (blue) which were subsequently analyzed for differential expression. For readability, the tree only shows the subtree of the NAM panel most relevant for the line selection.

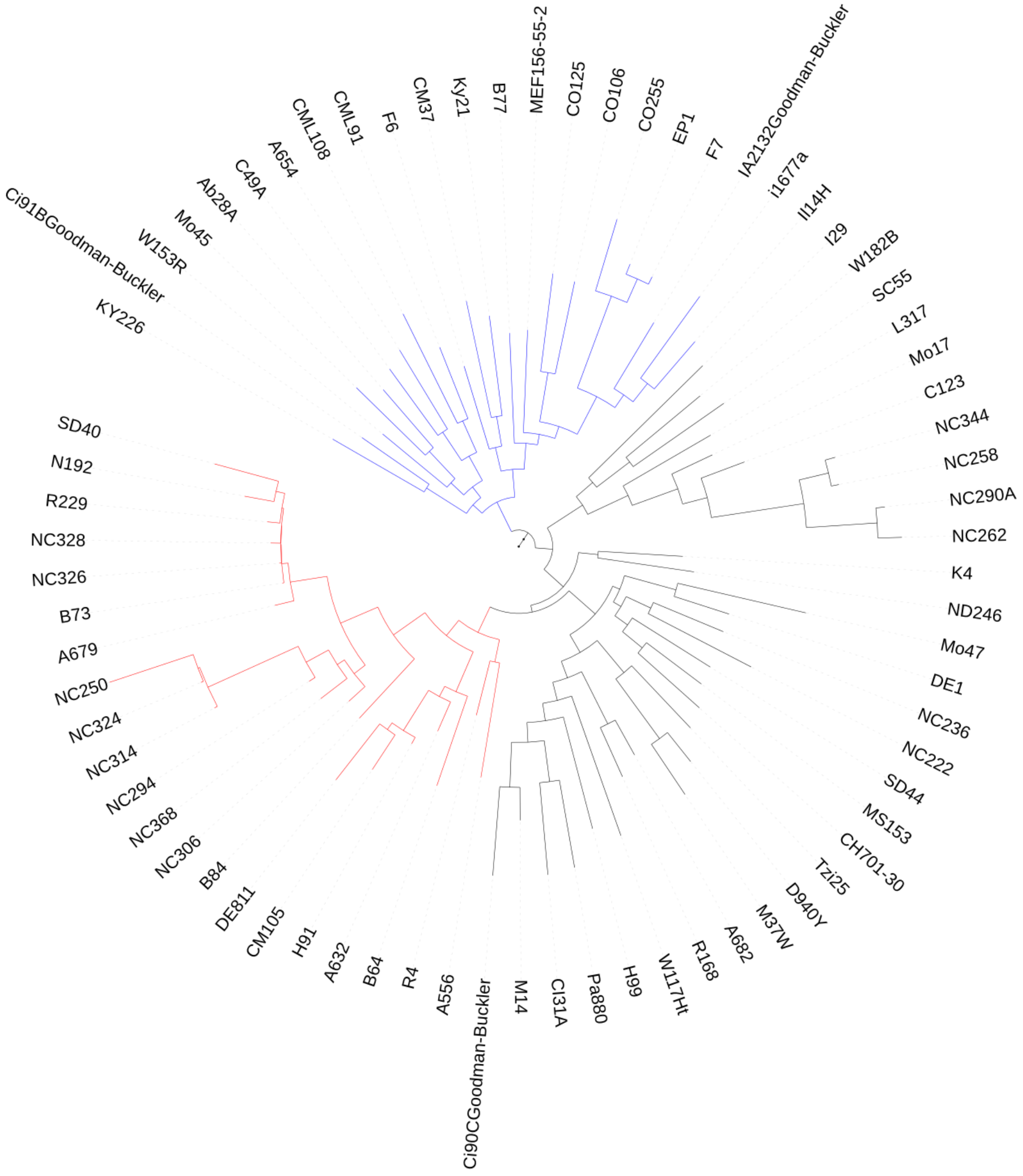
